# Contact-tracing in cultural evolution: a Bayesian mixture model to detect geographic areas of language contact

**DOI:** 10.1101/2021.03.31.437731

**Authors:** Peter Ranacher, Nico Neureiter, Rik van Gijn, Barbara Sonnenhauser, Anastasia Escher, Robert Weibel, Pieter Muysken, Balthasar Bickel

## Abstract

When speakers of two or more languages interact, they are likely to influence each other: contact leaves traces in the linguistic record, which in turn can reveal geographic areas of past human interaction and migration. However the complex, multi-dimensional nature of contact has hindered the development of a rigorous methodology for detecting its traces. Specifically, other factors may contribute to similarities between languages. Inheritance (a property is passed from an ancestor to several descendant languages), and universal preference (a property is universally preferred), may both overshadow contact signals. How can we find geographic contact areas in language data, while accounting for the confounding effects of inheritance and universal preference? We present `sBayes`, an algorithm for Bayesian clustering in the presence of confounding effects. The algorithm learns which similarities in a set of features are better accounted for by confounders, and which are due to contact effects. Contact areas are free to take any shape or size, but an explicit geographic prior ensures their spatial coherence. We test the clustering method on simulated data and apply it in two case studies to reveal language contact in South America and the Balkans. Our results are supported by —mostly qualitative— findings from previous studies. While we focus on the specific problem of language contact, the method can also be used to uncover other traces of shared history in cultural evolution, and more generally, to reveal latent spatial clusters in the presence of confounders.

## 1 Introduction

Speaker communities are rarely, if ever, completely isolated from each other. Communication between different communities requires finding a common language. This may lead to situations of bi- or multilingualism. Exposure to another language, especially if this is widespread within a community and takes place over a long period of time, can lead to horizontal transfer: the incorporation of words or structural features from one language into another. Although the importance of language contact for understanding the evolution of languages has been acknowledged already in the 19th century [57], modelling its effects remains a challenge in language data, and in patterns of cultural evolution more generally [16, 52, 10, 33, 46, 50, 5, 67].

Contact effects can take many shapes and sizes, and can be the result of a number of distinct processes. The most readily recognisable effects involve borrowing of forms (and functions) from one language to another. Commonly this involves the borrowing of lexicon (e.g. English borrowed the word *language* from French), but may also involve structural material, such as affixes or individual sounds (e.g. suffixes like *-able*, as in *readable* are borrowed from French).

When these types of contact effects spread from one language to another, it may lead languages spoken in a more or less contiguous area to become similar in their properties. The resulting areas of linguistic convergence are generally referred to as a linguistic area or *Sprachbund*. An example is the linguistic area of western and central Europe where languages tend to share several properties more commonly than in the adjacent regions of Asia, e.g. a system of definite and indefinite articles (English “the” vs “a”, Spanish “el/la” vs “un(a)”, Hungarian “a(z)” vs “egy”) [36]. Detecting such areas is challenging and problem-ridden [45, 10, 59, 5, 66], as they are the result of a number of complex historical processes that are difficult to reconstruct. How can we find geographical areas where languages have been in contact using empirical data and statistical inference?

A straightforward way of answering this question would be to look for shared features between geographically proximate languages. However, inferring contact from this alone ignores two important confounding effects that can also contribute to similarities between languages: inheritance and universal preference.

− Inheritance: Languages are transmitted from one generation to the next in an evolutionary process akin to the descent with modification that characterises biological evolution [14, 34]. In language, the modification stems from variation that each generation adds, mostly for signalling social identities. While this can lead to the split of a language into dialects and eventually into new languages, many properties persist and are inherited faithfully. As a result, languages may share a property just because they split from the same ancestral language and the property survived the split (or indeed several splits). An example is the inheritance of gender distinctions in many Indo-European languages (e.g. Italian, Russian, and Hindi).
− Universal preference: The structure of languages is shaped by universal aspects of how they are used for communication and thought, how they are processed in the brain, and how they are expressed with our speech and gesture systems. As a result,languages may share a property just because all languages tend to have it [4, 13, 31, 40, 44]. An example is the observation that virtually all languages have a formal means to distinguish questions from statements (e.g. intonation or a special word), with only very few exceptions [22].

Contact effects have generally been considered to be those (non-chance) similarities that are neither due to inheritance nor to universal preference. However, it is exceedingly difficult to attribute similarities categorically to contact, inheritance, or universal developments because the relevant processes interact in complex ways [5]. For example, a property that is universally preferred is also likely to be inherited when languages split, and to be borrowed in contact. Or, when languages are in contact over many generations, it is likely that they all tend to inherit the same properties. What is needed therefore, is a probabilistic way of estimating the relative contribution of each process.

In statistical terms, the task of finding contact areas can be described as clustering, i.e. finding groups of objects whose members share commonalities. However, naive clustering will simply group together languages with similar properties irrespective of the specific processes that have actually *made* them similar. Instead, we seek a method that infers the relative role of contact, as opposed to the other processes, in creating similarities between languages. Here we propose `sBayes`, a Bayesian mixture model which differentially weighs the respective contributions of contact and the confounding effects from inheritance and universal preference in accounting for the similarities between languages in space. While the model was primarily developed for linguistic data and we frame our discussion in terms of language contact, `sBayes` is applicable to a broader range of cultural evolution data. It is available as an open-source Python 3 package on https://github.com/derpetermann/sbayes, together with installation guidelines, a manual and case studies.

### Related work

The modern study of linguistic areas goes back to the early 20th century [8, 7, 62, 56]. The bulk of research since then has been qualitative in nature, but recently more quantitatively oriented approaches have been developed. We discuss the history of this strand of research in Section S1 of the Supporting Information. We conclude that a principled quantitative approach for finding contact areas is still missing, in particular one that takes into account both the process that leads to contact effects and the influence of confounding effects. A first approach to tackle this research gap was presented in [17], where a non-parametric Bayesian model was applied to reconstruct language areas. The approach distinguishes between areal and phylogenetic effects, but does not model universal preference as a confounder. A related idea was presented in [61], where an autologistic model together with family and neighbour graphs was used to assess the influence of inheritance and areality on cultural macroevolution in North America. The model does not itself infer areas, but instead assumes the spatial influence to happen within a fixed radius of 175 km. A somewhat different approach is proposed in [48]: based on prior knowledge a set of languages is assigned to a potential contact area—a “core”. Then a Naive Bayes classifier evaluates whether other languages belong to the core or to a control set, that is, languages unlikely to have been in contact with the core. The same authors also proposed a relaxed admixture model to detect language contact [11]. This mixture model locally detects borrowings between pairs of language, but does not reflect the possibility of larger contact areas.

Our method is inspired by these approaches, but, in contrast to them, it explicitly infers the assignment of languages to a contact area from the data: areas are allowed to take any possible shape and size and they are not constrained to a pre-defined sphere of influence. Instead, a geographic prior can be used to enforce spatial coherence and, thus, model the influence of geography. Moreover, the model controls for the two confounders of inheritance and universal preference, ensuring that only contact signals are picked up.

### Contact areas

We provide a data-driven definition of contact areas, which builds on linguistic features, that is, structural properties of language describing one aspect of cross-linguistic diversity (as e.g. found in [23]). Consider a set of languages *L* = {*l*_1_, *l*_2_, *…*}, for which we study the feature *f*_palatal_, the presence and absence of palatal nasals, an item of the phonological inventory. Suppose further that there is an area *A* where palatal nasals are present in all languages, while they are commonly absent everywhere else. Universal preference fails to explain why languages in *A* have palatal nasals. We might conclude that we found evidence of some form of shared history, either due to inheritance or contact—making *A* an *area of shared history*. Clearly, this conclusion is weak: it builds on a single source of evidence and neglects chance, which becomes apparent once the distribution of a feature is less clear-cut (Fig. 1a). Inside the dashed-line polygon (*A*), languages are roughly twice as likely to have palatal nasals than outside *A*. Languages inside the polygon are similar and universal preference does not explain why. And yet, it seems arbitrary to conclude that *A* shows shared history. All the same it seems equally arbitrary to simply disregard the similarity in *A* altogether.

**Figure 1:**
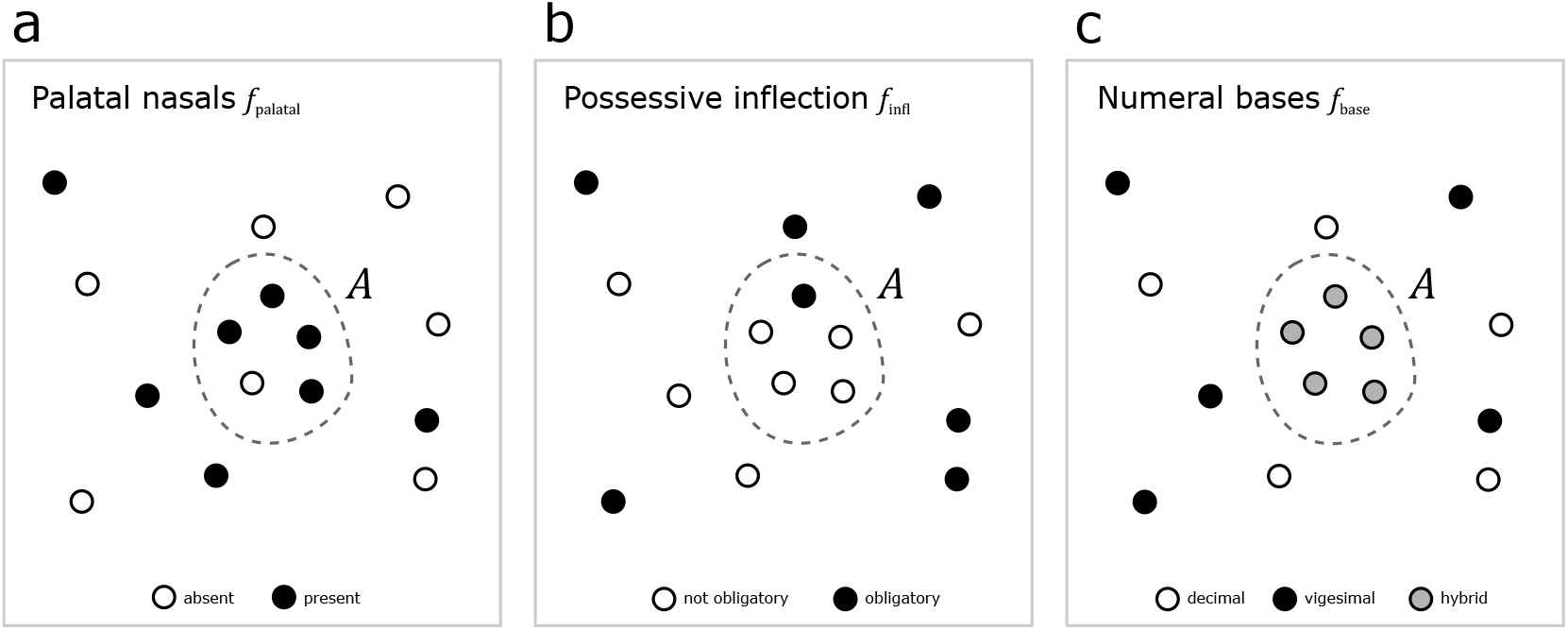
Area of shared history. In the area *A*, features *f*_palatal_, *f*_infl_ and *f*_base_ (dashed-line polygon) follow a distribution with low entropy, which differs from the distribution outside of *A*. Note that the features only serve illustration here; for definitions and actual distributions, see the World Atlas of Language Structure [23].

A standard response is to consider additional, independent features that reinforce or weaken the similarities observed for a single feature. Suppose we also study the grammatical feature *f*_infl_, the presence and absence of obligatory possessive inflection, and the lexical feature *f*_base_, the type of base system used for expressing numerals. For most languages in *A*, possessive inflection is not obligatory (Fig. 1b). Moreover, all languages in *A* use the same hybrid vigesimal-decimal base system (Fig. 1c). Each additional feature reinforces the signal observed for palatal nasals. More formally, across all three features languages in *A* have low entropy, i.e. they are similar and thus predictable, and differ from the confounder, i.e. they cannot be explained by universal preference.

#### Definition 1.1.

In an area of shared history *A*, independent features *f*_1_, *f*_2_, … *f*_n_ follow a distribution with low entropy, which differs from the distribution expected from the confounding effect of universal preference.

Definition 1.1 ensures that a random accumulation of universally preferred features is not mistaken for shared history. The definition is largely impartial to the argument that preferred features are also more likely inherited and shared. For example, subject-before-object orders are universally preferred over object-before-subject orders [51, 21] but the global distribution still shows geographic structure: some areas, such as Eurasia, Africa, or Papua New Guinea, show an even stronger preference than the worldwide norm. Thus, even strongly preferred patterns can provide evidence for an area.

Definition 1.1 separates unspecified shared history from universal preference, but does not distinguish between contact and inheritance: features in *A* could have been passed on from neighbours or they could have been inherited. We adapt the definition to account for the confounding effect of inheritance and, thus, isolate similarities due to contact.

To approach this issue, let us assume, as an example, that most languages are related to others and belong to a language family *ϕ* ∈ Φ, where Φ is the set of all language families. Languages in Fig. 2 belong to either family *ϕ*_blue_ or *ϕ*_red_. Let us further assume that there are two areas *A* and *Z* that both contain four languages from *ϕ*_red_ and one from *ϕ*_blue_. In both areas the entropy of each linguistic feature is lower in *A* and *Z* than is the case outside, in the entire set of languages. However, all languages in *A* have features that are also common in *ϕ*_red_ (Fig. 2a), while they are relatively rare for languages in *Z* (Fig. 2b). Inheritance explains the similarity in *A*, but it fails to explain the similarity in *Z*. Thus, *Z* is a contact area, whereas *A* is not. With this, we arrive at the final definition of a contact area:

**Figure 2:**
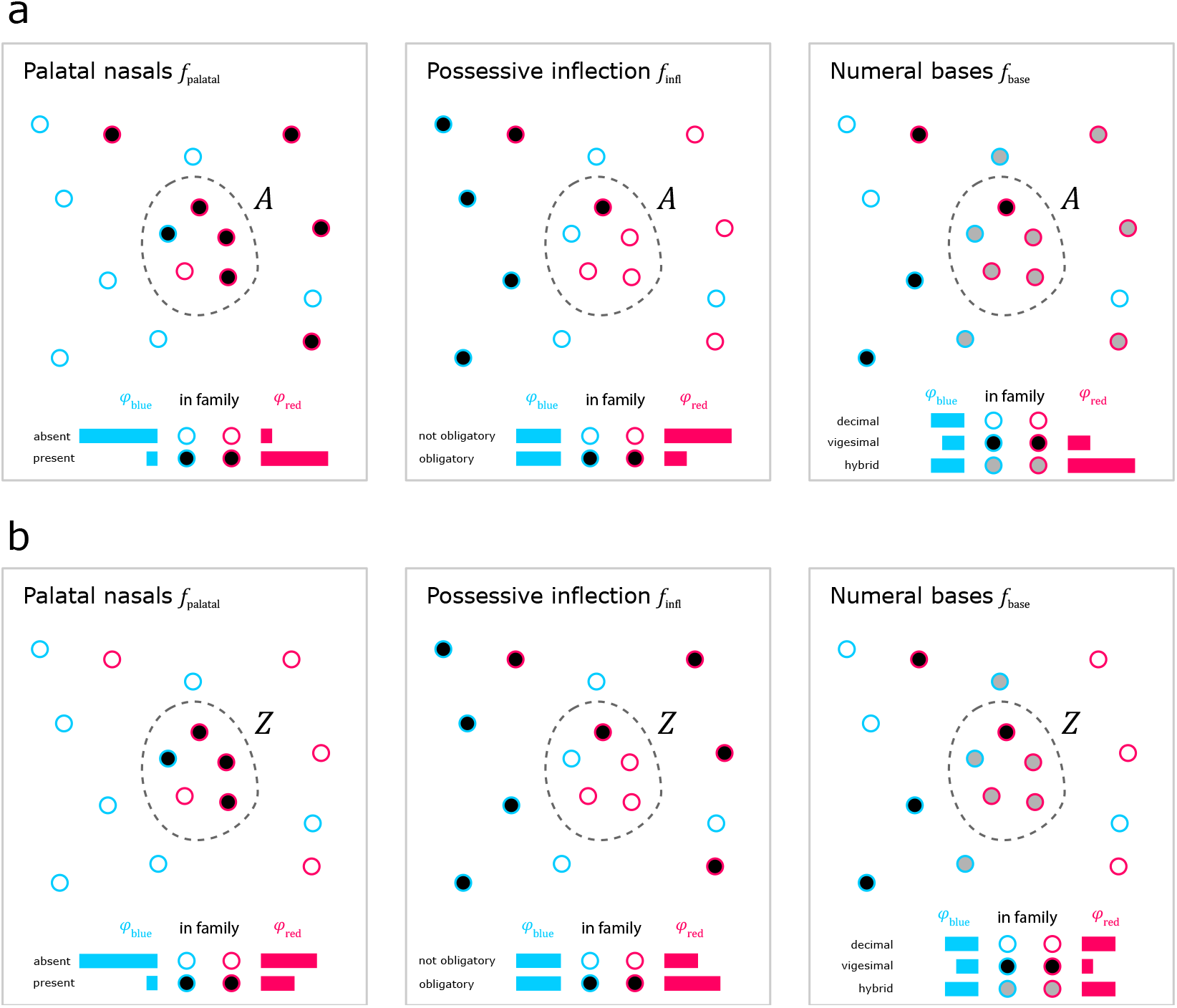
Contact areas: In the areas *A* and *Z* (dashed-line polygons) features *f*_palatal_, *f*_infl_ and *f*_base_ follow a distribution with low entropy, which differs from the distribution outside the polygons. The blue and red horizontal bars show how common a feature is in each family. (a) The distribution in *A* largely matches the distribution in family *ϕ*_red_. *A* can be explained by inheritance and is not a contact area. (b) The distribution in *Z* does not match the distribution in *ϕ*_red_. Inheritance fails to account for the similarity in *Z*, which leaves contact as the remaining explanation: *Z* is a contact area.

#### Definition 1.2.

In a contact area *Z*, independent features *f*_1_, *f*_2_, … *f*_n_ follow a distribution with low entropy, which differs from the distribution expected from the confounding effect of universal preference. Moreover, the distribution in *Z* also differs from the distribution in families Φ and, thus, cannot be explained by the confounding effect of inheritance.

Based on definition 1.2, we introduce `sBayes`, an algorithm to find contact areas on the basis of language data.

## 2 Materials and Methods

`sBayes` requires features to be categorical. A feature *f* is assumed to have *k* discrete, mutually exclusive states

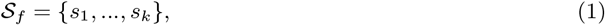

where *𝒮*_*f*_ is the set of states and *s*_1_, *…, s*_*k*_ are the state labels. For example, palatal nasals have two states, they can be present or absent: *𝒮*_palatal_ = {present, absent}. Ideally, each state is self-contained and carries explicit information about shared history, which is the case for *𝒮*_infl_ = {decimal, hybrid, vigesimal}, but less so for *𝒮*_infl_ = {decimal, hybrid, other}, since the state *other* does not refer to a well-defined base system.

### Likelihood

The model aims to identify effects that predict why feature *f* in language *l* has state *s*. `sBayes` proposes three effects and defines a likelihood function for each:

− likelihood for universal preference (*P*_universal_): the state is universally preferred.
− likelihood for inheritance (*P*_inherit_): the language belongs to family *ϕ*(*l*) and the state was inherited from related ancestral languages in the family.
− likelihood for contact (*P*_contact_): the language belongs to area *Z*(*l*) and the state was adopted through contact in the area.

`sBayes` models each feature as coming from a distribution that is a weighted mixture of universal preference, inheritance and contact. The unknown weights — *w*_universal_, *w*_inherit_ and *w*_contact_ — quantify the contribution of each of these three effects. For a single language *l*, a feature *f* and a state *s*, this yields the following mixture model:

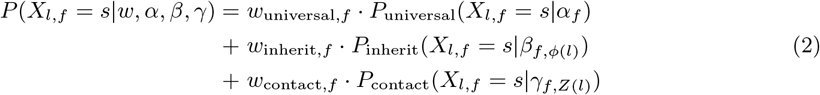

The mixture components — *P*_universal_, *P*_inherit_ and *P*_contact_ — are categorical distributions parameterised by probability vectors *α*_*f*_, *β*_*f,ϕ*(*l*)_ and *γ*_*f,Z*(*l*)_. That is, the probability of observing state *s* in feature *f* is *α*_*f,s*_ if it is the result of universal preference, *β*_*f,ϕ*(*l*),*s*_ if it was inherited in family *ϕ*(*l*) and *γ*_*f,Z*(*l*),*s*_ if it was acquired through contact in area *Z*(*l*). While the assignment of languages to families is fixed, the assignment of languages to areas is inferred from the data. `sBayes` allows for multiple contact areas *Ƶ* = {*Z*_1_, *Z*_2_, *…*}, each with their own set of areal probability vectors. A detailed explanation of all mixture components together with examples can be found in section S2 of the Supporting Information. The weights *w*_*f*_ = [*w*_universal,*f*_, *w*_inherit,*f*_, *w*_contact,*f*_] model the influence of each component on a feature:

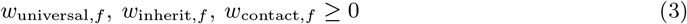

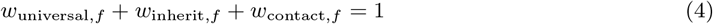

For languages not assigned to a contact area, the contact weight is set to zero and the other weights are re-normalised accordingly.

The mixture model combines the likelihood for universal preference, inheritance, and contact and their weights across all languages. The model has parameters Θ = {*Ƶ, α, β, γ, w*}, which are evaluated against the data *D*, that is, the states of all features in all languages. The likelihood of the whole model for the given data is the joint probability of states *s* over languages *l* ∈ *L* and features *f* ∈ *F*, given Θ:

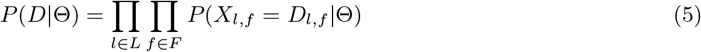

### Model intuition

`sBayes` preferentially samples areas with high likelihood values. This is the case if estimates for the areal probability vector, *γ*_*f,Z*(*l*)_,

‐ fit the data
‐ have low entropy
‐ differ from the probability vectors of the confounders

Fig. 3a illustrates how `sBayes` evaluates evidence for contact for a single feature with two states *A* (blue) and *B* (yellow). The distribution of the feature in the proposed area *Z* has low entropy (blue and yellow columns) and differs from the distribution of the two confounders – universal preference (solid black line) and inheritance (dashed black line). This pulls the weights vector (pink star) towards contact. Fig. 3b shows that given the same confounding effect the likelihood increases with increasing entropy in area *Z*. `sBayes` avoids areas where universal preference and inheritance explain the similarity in the data equally well or even better than contact, but instead picks up areas for which the confounders do not provide an adequate explanation, given that their entropy is low.

**Figure 3:**
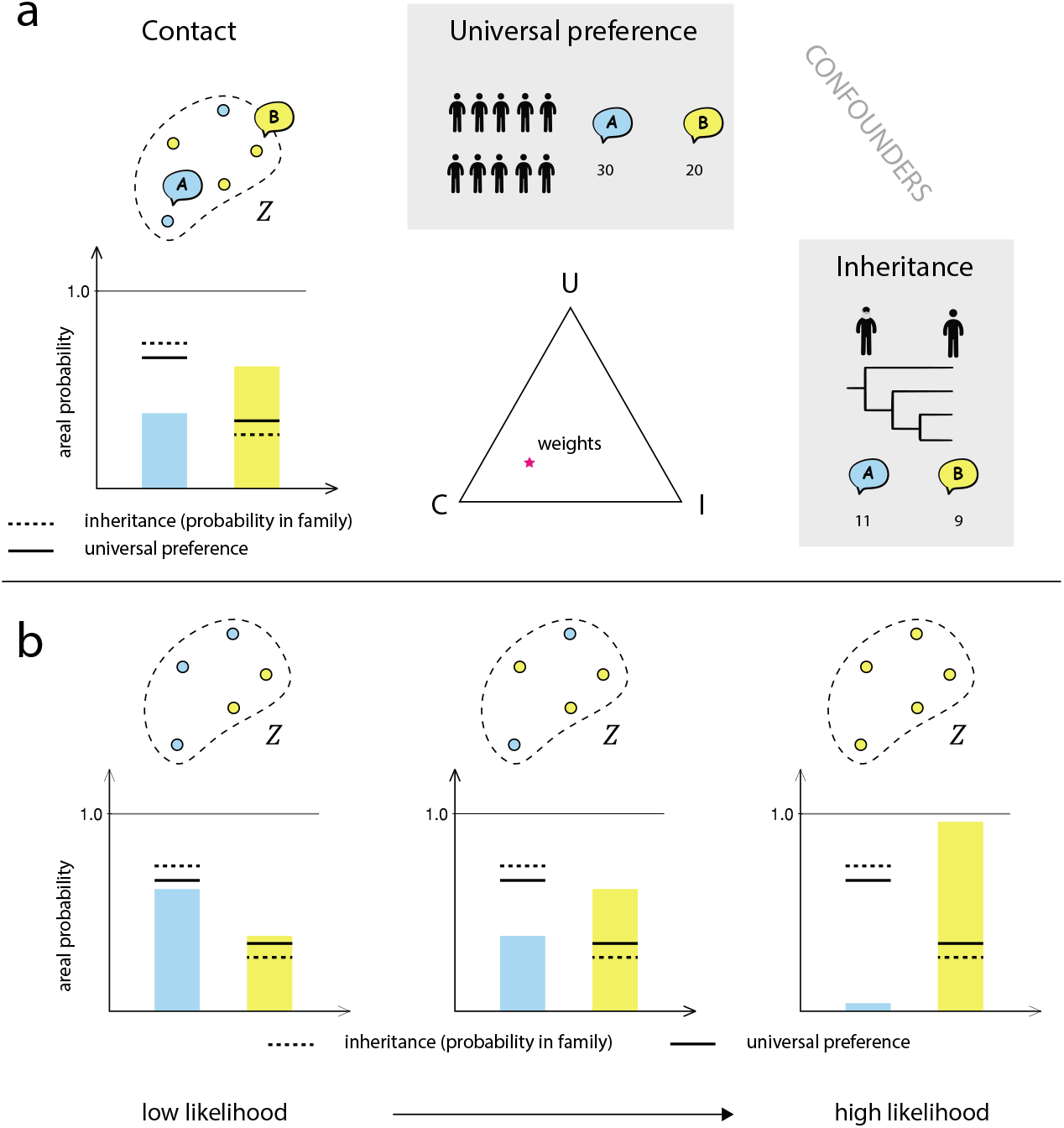
(a) The languages in area *Z* are explained better by contact than universal preference and inheritance. The weights vector (pink star) leans towards contact. (b) The likelihood of the model is highest when the areal probability vector has low entropy (i.e., features in *Z* are similar) and when it differs from the confounders.

### Prior

In `sBayes`, priors must be defined for the mixture weights and the probability vectors of the categorical distributions on the one hand, and the assignment of languages to areas on the other. `sBayes` uses Dirichlet priors for the weights and the probability vectors and purpose-built geo-priors for the assignment of languages to areas:

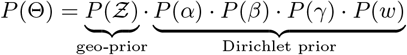

Both the weights *w*_*f*_ and the vectors *α*_*f*_, *β*_*f*_, *γ*_*f*_ parameterise a categorical distribution: they are bounded between [0, 1] and sum to 1, which motivates the use of a Dirichlet prior. The default prior is uniform:

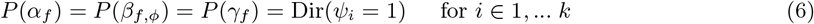

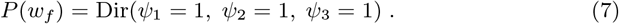

In other words, any of the *k* states and any of the three weights are equally likely *a priori*. While this invariance seems reasonable for the weights, it might not always be appropriate for the probability vectors: *α*_*f*_ allows the model to learn which states are universally preferred, and *β*_*f,ϕ*_ which states are inherited in family *ϕ*. The more a state is preferred universally or in a family the less likely a similar occurrence in *Z* is regarded as evidence for contact. However, what is rare in our sample (i.e., our study area) might be abundant outside and vice versa.

In an ideal setting `sBayes` would be applied to a global sample of languages, making it possible to infer universal preferences directly from the data. When this is not possible, preference may be incorporated in the form of an empirical prior. The prior allows us to express specific knowledge about universal preferences before seeing the data. In the Dirichlet distribution, the parameters *ψ*_*i*_ can be thought of as pseudocounts for each of the *k* states, which can reflect prior knowledge:

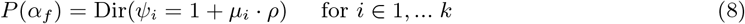

In Equation 8, *µ*_*i*_ is the prior probability of state *i* and defines the mean of the prior distribution, while *ρ* gives the precision or inverse variance. A large *ρ* implies a strong prior with low variance. An informative prior for inheritance in family *ϕ*, i.e. *P* (*β*_*f,ϕ*_), is defined analogously. In Section S3.1 of the Supporting Information, we illustrate how a biased sample might lead to biased estimates for universal preference and we provide an example for an empirically informed prior.

Each language *l* is geographically situated: it has a spatial location, that is, a unique point in geographical space (if we assume languages to be represented by their centre of gravity). The geo-prior models the *a priori* probability of languages in an area to be in contact, given their spatial locations. `sBayes` employs two types of geo-priors:

− a *uniform* geo-prior
− a *cost-based* geo-prior.

The *uniform* geo-prior assumes all areas to be equally likely, irrespective of their spatial locations, whereas the *cost-based* geo-prior builds on the assumption that close languages are more likely to be in contact than distant ones. Distance is modelled as a cost function *C*, which assigns a non-negative value *c*_*i,j*_ to each pair of locations *i* and *j*. Costs can be expressed by the Euclidean distance, great-circle distance, hiking effort, travel times or any other meaningful property quantifying the effort to traverse geographical space. Since costs are used to delineate contact areas, they are assumed to be symmetric, hence *c*_*i,j*_ = *c*_*j,i*_. For cost functions where this is not immediately satisfied the cost values can be made symmetric, e.g. by averaging the original costs.

`sBayes` finds the minimum spanning tree *T*_*Z*_, which connects all languages in *Z* with the minimum possible costs (red and blue lines in Fig. S1b, Supporting Information). *T*_*Z*_ quantifies the *least* effort necessary for speakers in *Z* to *physically meet* given the particular cost function used. We define the cost of an area as the average cost over all edges in the minimum spanning tree

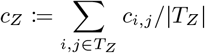

and let the prior probability decrease exponentially as the average cost increases:

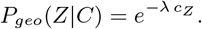

The parameter *λ* defines the rate at which the probability decreases. A large *λ* results in a strong geo-prior: distant languages with high costs have very low prior probability to allow for contact. When *λ* is small, the exponential function becomes flat and the geo-prior approaches a uniform distribution (Fig. S1b, Supporting Information). Using the average (rather than the sum) to define the cost of an area, ensures that this prior is agnostic to the number of languages in *Z*. The geo-prior not only expresses our belief that spatial proximity leads to contact, but also our confidence in the present-day locations of languages, which might have been different just a few hundred years ago.

In addition to the geo-prior, there are two implicit parameters relating to the prior probability of contact areas: the size of an area in terms of number of languages, *m*, and the number of areas, *n*. The prior for *m* is discussed in section S3 of the Supporting Information. There is no prior for *n*. Instead, we run the model iteratively, increase the number of areas per run and compare the performance across *n* in postprocessing (see below).

### Posterior

The posterior of the model is proportional to the likelihood times the prior:

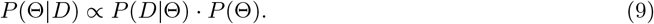

Section S5 of the Supporting Information, explains how `sBayes` samples from the posterior distribution *P* (Θ|*D*) to identify potential contact areas. `sBayes` employs a Markov chain Monte Carlo (MCMC) sampler with two types of proposal distributions: a Dirichlet proposal distribution for weights and probability vectors and a discrete, spatially informed proposal distribution for areas.

### Number of areas

With more areas `sBayes` will find it easier to explain the variance in the data. However, each area requires additional parameters, resulting in a more complex model and higher uncertainty in the posterior. `sBayes` employs the *deviance information criterion* (DIC) to find a balance between fit and complexity. The DIC estimates the effective number of parameters from the uncertainty in the posterior and uses it to penalise the goodness of fit [58]. We run `sBayes` iteratively increasing the number of areas, *n*, and evaluate the DIC for each run. The most suitable *n* is where the DIC levels off, such that adding more areas does not improve the penalised goodness of fit. The DIC has been found to outperform competing approaches for identifying the optimal number of clusters in a comparable Bayesian clustering procedure [29]. We show that the DIC correctly reports the true number of areas in simulated data (Section S7 of the Supporting Information). However, the DIC is not part of the core methodology and can be replaced with other model selection criteria, e.g. the WAIC [69] or PSIS-LOO [68].

Once a suitable *n* has been identified, areas are ranked according to their relative posterior probability in post-processing (see section S4, Supporting Information).

## 3 Results

For all experiments we ran `sBayes` with 3 million iterations, of which the first 20% were discarded as burn-in. We retained 10,000 samples from the posterior and used Tracer [54] to assess the effective sample size and convergence.

### Simulation study

Before applying `sBayes` to real-world data, we performed a simulation study to verify that the algorithm correctly samples from the posterior distribution under model assumptions. We assigned ∼ 950 languages to random locations in space and simulated 30 features for each to model universal preference. All features were generated according to a categorical distribution with two, three, or four states. We carried out four experiments:

− *Experiment 1* correctly identified contact areas differing in shape, size and strength of the signal.
− *Experiment 2* distinguished between similarity due to inheritance and due to horizontal transfer, separating contact effects from inheritance in a family.
− *Experiment 3* correctly estimated the number of contact areas.
− *Experiment 4* used empirically informed priors to robustly infer contact areas even for small and biased samples.

Experiment 2 will be explained in more detail below. All remaining simulation experiments can be found in Section S7 of the Supporting Information. Experiment 2 demonstrates that `sBayes` distinguishes between similarities due to inheritance and those due to contact. We assigned some of the simulated languages to a common language family and some to a contact area. We simulated shared ancestry in the family and contact in the area with different categorical distributions. The entropy for inheritance was set to be lower than that of contact, i.e., the signal for shared ancestry was assumed to be stronger. Finally, we simulated weights controlling the influence of each effect. Then, `sBayes` was run with two different setups. In the first setup, the information about common ancestry was not passed to the algorithm. `sBayes` incorrectly attributes the similarity in the family to contact. Assuming a single contact area (*n* = 1), the posterior of *Z*_1_ overlaps with the simulated language family, but misses out on the weaker simulated contact area (Fig. S4a, Supporting Information). In the second setup, inheritance was modelled at the family level and passed to the algorithm. Now, `sBayes` was able to learn that the similarity in the family was due to inheritance. The posterior correctly returns the simulated contact area (Fig. S4b, Supporting Information).

`sBayes` not only finds contact areas, but also infers the influence of each feature to delineate them. Fig. 4 shows the simulated values for universal preference, inheritance in the family, and contact in *Z*_1_ (pink star) for three features, and their inferred posterior distribution (heat map ranging from yellow to dark blue). Feature f6 (Fig. 4a) is strongly shared in *Z*_1_; both the simulated and inferred weights lean towards contact. In the area, most languages have either state 1 or 2. In the family, state 0 is preferred. Universally, there is no preference for either state. Feature f3 (Fig. 4b) is both inherited in the family and shared in the area. The simulated and inferred weights lie between contact and inheritance. Feature f4 (Fig. 4c) is indecisive. The simulated weights lie in the centre, the inferred estimates scatter across the entire probability simplex.

**Figure 4:**
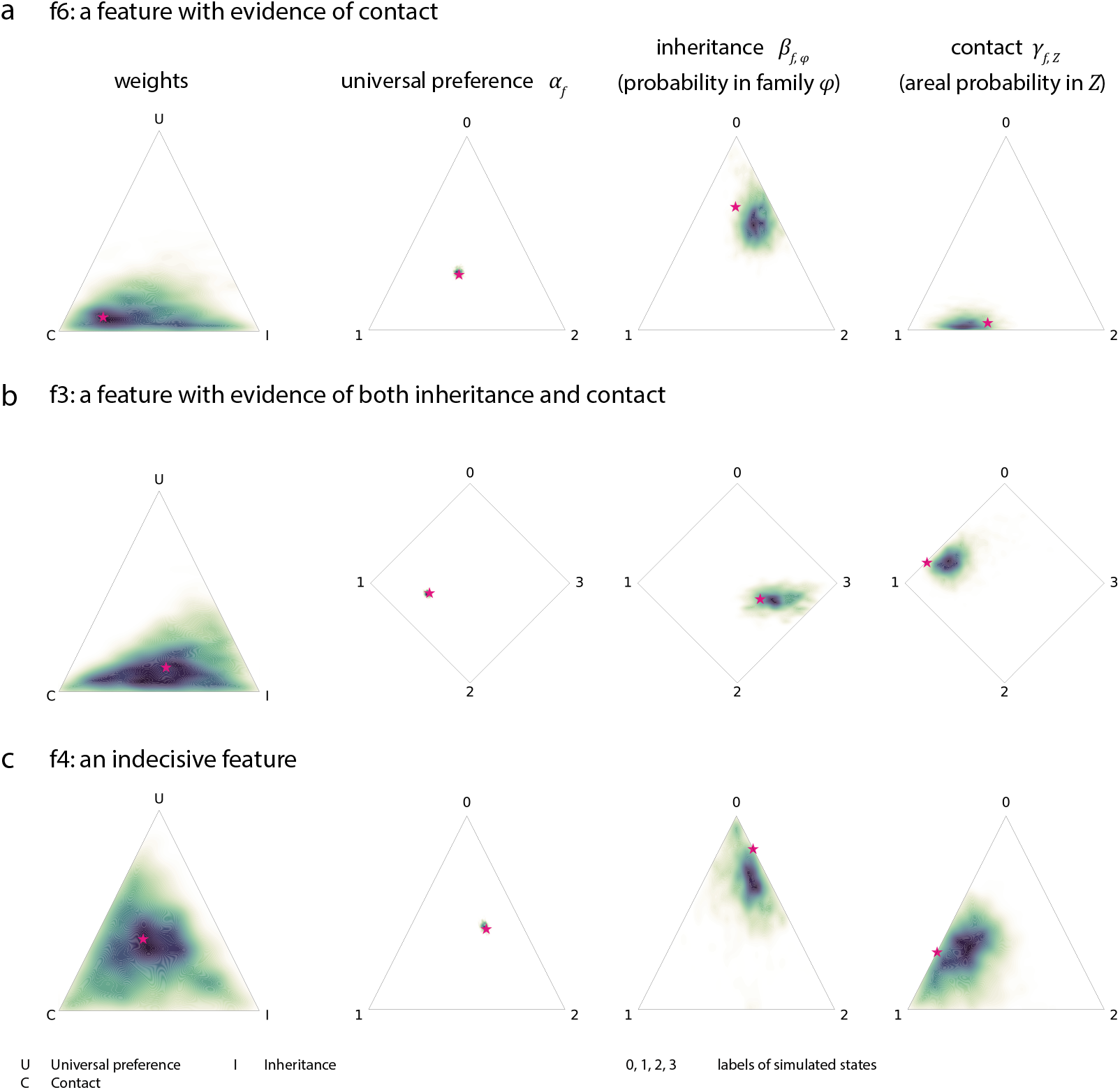
Simulated and reconstructed weights and states (U: universal preference, I: probability of inheritance in family *ϕ* and C: probability of a contact effect in area *Z*) for three features (f6, f3 and f4). The heat map shows the probability density of the posterior distribution. The pink star marks the ground truth value, i.e., the simulated weights or states. Feature f6 provides evidence of contact (a), f3 of inheritance and contact (b), f4 is indecisive (c).

### Case Study: Western South America

Western South America is characterised by extreme genealogical diversity. At the same time, the region contains a number of putative linguistic areas, where languages share a number of grammatical characteristics as a result of contact [24]. These linguistic areas exist against the background of a proposed major split between Amazonian lowland languages on the one hand and Andean highland languages on the other [9, 19, 20, 60].

The data set used for the case study consists of 100 languages presently spoken in the western Amazon basin and adjacent Andean highlands (Fig. S7, Supporting Information). The 100 languages were coded for 36 features of grammar (Table S2, Supporting Information). The prior for universal preference was derived from a stratified global sample (86 languages from different language families spread uniformly over the globe). The mean of the Dirichlet prior was set equal to the mean of the stratified sample. The precision was set to 10, yielding a weakly informative prior. Inheritance was modelled for families with at least five members: Arawak, Panoan, Quechuan, Tacanan, Tucanoan and Tupian. A prior for each family was derived from 37 languages outside the sample analogously to the universal prior, except for Tacanan, for which all (known) members were in the sample and a uniform prior was used instead. The geo-prior was set to be uniform. Fig. 5 shows the results of the experiment. Language families are shown by shaded areas, contact areas by coloured lines. We ran the analysis iteratively, increasing the number of areas per run. The DIC starts to level off for *n* = 3, suggesting three salient contact areas in the data (Fig. S8, Supporting Information).

**Figure 5:**
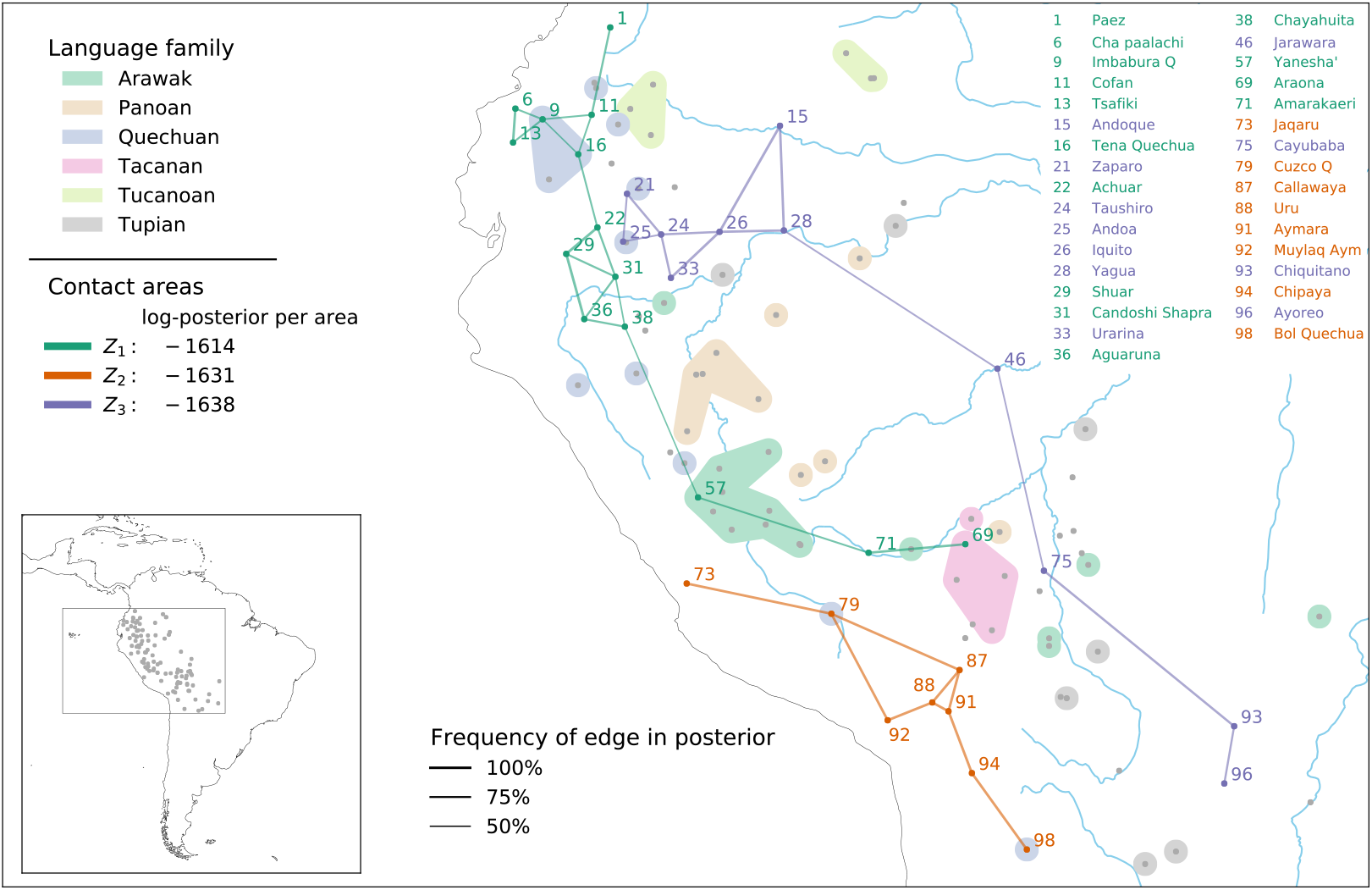
Contact areas in Western South America: The posterior distribution consists of contact areas *Z*_1_, *Z*_2_ and *Z*_3_ (connected by green, orange and purple lines), ordered by posterior probability. The grey dots indicate the spatial locations of all languages in the sample, the shaded areas represent the six main language families. Languages in each area are connected with a Gabriel graph [47], line thickness corresponds to the frequency of an edge in the posterior (how often are two adjacent languages together in the same area?).

The northern part of *Z*_1_ has likely been an area of interethnic interaction for a long time, connected to the sphere of influence of the Chibcha family [12, 15, 1], and smaller-scale interactions into the lowlands (e.g. [72, 65, 41]). Of the geographically more remote languages Yanesha’ and Araona (or more generally the Tacanan family) have known historical contact relations with Quechuan languages [2]. Moreover,it has been observed that Yanesha’ shares features with the northern cluster [72]. Amarakaeri is a relatively recent arrival in the foothills, with looser ties to the Incas [1, 2]. The contact features contributing most to the northern area are generally associated with Andean languages [20, 60] (Fig. S10, Supporting Information).

Area *Z*_2_ corresponds to a well-known case of intensive language contact between Aymaran and Quechuan languages and, more peripherally, the Uru-Chipaya family [60, 2]. The most likely contact features in our analysis correspond to known Andean features, mostly phonological (Fig. S11, Supporting Information). Areas *Z*_1_ and *Z*_2_ roughly correspond to the northern and southern central Andes, respectively, (with some incursions into the lowlands). This is consistent with recent results [48, 64], which suggest that the Andes consist of “two distinguishable but interlocking linguistic areas, one northern and one southern” [64].

Amazonian-based area *Z*_3_ is spread over a large territory, which may be due to the fact that Amazonian languages, generally speaking, lack a number of features that are characteristic for Andean languages. This is corroborated by the most contributing features which mark the absence of typical Andean characteristics (Fig. S12, Supporting Information). The densest part of this area, however, may be connected to the idea of a larger trade area around the Marañon River [71, 64], ultimately connected to the north-western part of a vast trade area [25, 38]. A contributing reason for the connection between the northern and southern clusters of area *Z*_3_ may be the fact that the two largest families of the continent, Arawak and Tupian, have branches that extend into the northwest Amazon as well as the Madeira-Guaporé-Mamoré area in the south.

### Case Study: Balkans

The Balkans data set consists of 30 languages and dialects situated within and outside the geographical boundaries of the Balkan peninsula: Albanian, Macedonian, Bulgarian, Torlak, Aegean Slavic, Bosnian-Croatian-Montenegrin-Serbian, Aromanian, Istroromanian, Romanian of Romania and Moldova, and Balkan Turkish (Fig. S13, Supporting Information). With the exception of Turkish, they all belong to the Indo-European family. The 30 varieties were coded for 47 features from various linguistic domains (see Table S3, Supporting Information). Inheritance was modelled at the sub-clade level for Albanian, Greek, Romance and Slavic dialects and at the family level for Turkic. We used a stratified sample of 19 European languages to model a prior for universal preference (or, in this case, Standard Average European preference). The mean of the Dirichlet prior was set equal to the mean of the European sample. The precision was set to 10, resulting in a weakly informative prior. Analogously, we collected 23 languages outside the sample to derive empirically informed priors for all sub-clades, except for Albanian, for which all members were in the sample and a uniform prior was used instead. The geo-prior was set to be uniform. Fig. 6 shows the results of the experiment. Language families are shown as shaded areas, contact areas by coloured lines. We ran the analysis iteratively, increasing the number of areas per run. The DIC levels off for *n* = 3, after which it increases sharply, suggesting three areas in the data (Fig. S15, Supporting Information).

**Figure 6:**
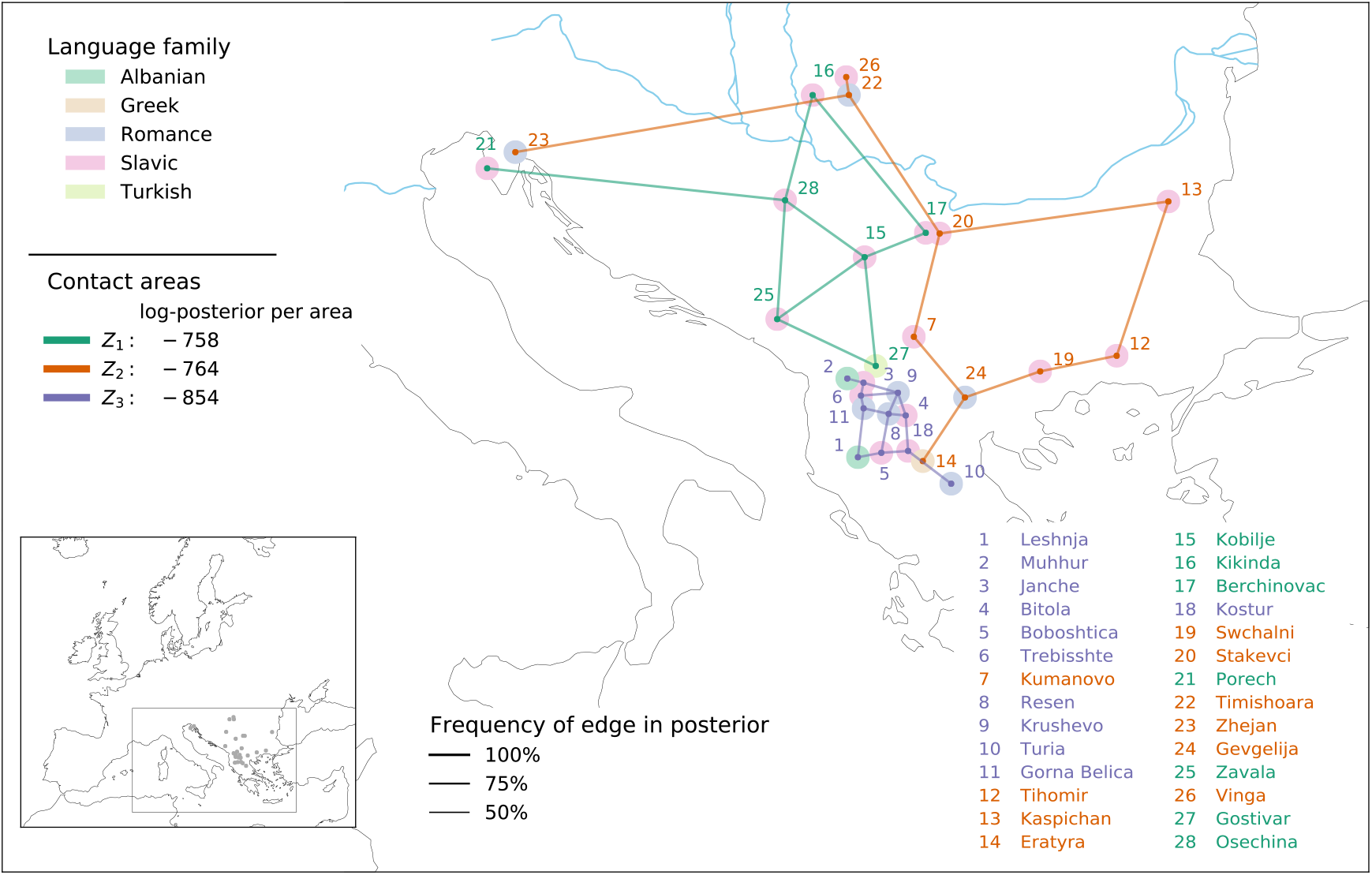
Contact areas in the Balkans: The posterior distribution consists of contact areas *Z*_1_, *Z*_2_ and *Z*_3_ (connected by green, orange and purple lines) ordered by posterior probability. The shaded circles represent the sub-clades and language families. Languages in an area are connected with a Gabriel graph, line thickness corresponds to the frequency of an edge in the posterior (how often are two adjacent languages together in the same area?).

The dialects in the sample share a common history that differs from both Standard European preference and the family probability in each of the sub-clades. For *n* = 1, all dialects are assigned to one single area—except for the two Albanian dialects of Leshnja and Muhhur, and the Turkish dialect of Gostivar, which are still reasonably well explained by inheritance in the Albanian sub-clade and in the Turkic family, respectively (Fig. S14, Supporting Information). This single Balkan area divides into three salient areas (Fig. 6), now also including the above three dialects.

Area *Z*_1_ joins different varieties of the southwestern part of the Serbo-Croatian dialect continuum. The area is distinct within the Slavic branch and—as indicated in Fig. S17, Supporting Information—is defined through a lack of features that would traditionally be expected in the Balkan Sprachbund [37, 3]. Dialects in *Z*_1_ had almost no contact with Albanian and Romance/Aromanian and were not exposed to the processes of language convergence observed in areas *Z*_2_ and *Z*_3_. The fact that Gostivar Turkish belongs to *Z*_1_ indicates that it has converged with these varieties in certain respects.

Area *Z*_2_ includes the Greek variety of Eratyra, the Meglenoromanian variety of Gevgelia, all Bulgarian dialects, the Romance varieties to the north of the Danube, i.e. Slavic dialects spoken in the Aegean, Slavic dialects in a Romance surrounding and Romance dialects in a Slavic surrounding. The area shows Romance–Slavic and Slavic–Greek contacts. Interestingly, some of the defining features (F30, F33, F38; Fig. S18, Supporting Information) are characteristic of Albanian dialects and are also shared in *Z*_3_, suggesting contact between the two areas. This is also reflected when running `sBayes` with *n* = 2, in which case *Z*_2_ and *Z*_3_ are merged into a single large area.

Area *Z*_3_ comprises all Albanian and Aromanian, as well as the western Macedonian Slavic dialects. The area shows intense contact and multilingualism, characterised by a set of properties for which a contact explanation is the most probable one (Fig. S19, Supporting Information). This corresponds to what is known from traditional studies on the Balkan area, which identify the area around lake Ohrid and along the border between today’s Albania and North Macedonia as the centre of areal innovations [32, 42]. Overall, Slavic varieties partake in all three areas. `sBayes` clearly divides West South Slavic and East South Slavic. The former constitutes an area mainly by its divergence within the Slavic branch as a result of dialect contacts. Whether these varieties are also part of another convergence zone, e.g. with the languages of the Austro-Hungarian Empire, remains to be investigated with additional data. East South Slavic is affected by different contact situations: with Romance and Greek in *Z*_2_, with Romance and Albanian in *Z*_3_. In this way, a historical interpretation of the three areas seems possible: *Z*_1_ is the oldest area of internal South Slavic dialect contact (Turkish joining later), *Z*_2_ shows contact fostered by the Byzantine Empire, while *Z*_3_ reflects contact triggered within the Ottoman Empire. In any case, contact with Albanian emerges as the crucial element responsible for the specific Balkan convergence processes in *Z*_3_. In sum, the three areas largely confirm what is known from traditional studies, albeit on a strictly empirical basis and disclosing the relevant premises.

## 4 Discussion

We presented `sBayes`, a Bayesian clustering algorithm to identify areas with similar entities while accounting for confounders. Specifically, we tailored the approach to language data and identified areas of language contact, while accounting for universal preference and inheritance. We tested the approach on simulated data and performed two case studies on real-world language data in South America and on the Balkans.

### Model assumptions and diagnostics

Our model assumes that contact leaves behind traces in extant languages in the form of areas, which emerge once the more salient traces of confounding effects have been properly accounted for. Specifically, the mixture model assumes that each feature in each language is explained probabilistically by three effects: universal preference, inheritance in a family, and contact in an area. `sBayes` iteratively proposes areas and evaluates them against the data. Areas have a high likelihood for contact if they comprise similar features which cannot be equally well explained by universal preference and inheritance. There are no assumptions about any of the properties of contact areas, such as their shape, size or number, whether they comprise close or distant languages, or cover contiguous or disconnected regions in space. The algorithm learns these properties from the data, potentially guided by informative (geographical) priors. Likewise, `sBayes` is agnostic to features and their relationship to borrowing. A priori, all features are treated as equal and independent evidence. Proposing and evaluating contact areas in turn, the algorithm learns which features are better explained by each of the three effects. In this sense, the analysis is data-driven: only sufficient, informative, and independent features provide a robust statistical signal to delineate contact areas.

`sBayes` does not replace expert knowledge in defining the features, the confounders (e.g. the families), the priors, and the spatial locations and in interpreting the results in an anthropological and historical context. In the absence of salient contact areas in the data, `sBayes` might group together outlier languages that are poorly explained by either of the confounders. `sBayes` provides statistics and measures to detect such spurious areas in the posterior. MCMC diagnostics assess whether sampling has converged to a stable, stationary distribution and whether the posterior contains sufficient independent samples (Section S5, Supporting Information). Measures of model fit evaluate the evidence for contact in the posterior. Spurious areas have a high entropy and a low likelihood, resulting in a high DIC. Priors account for biased data and enforce spatial plausibility. However, statistics and priors can only address the internal validity of the model. Spurious areas can still arise because of misspecified confounders, e.g., the algorithm returns a language family that was not included in the model, or redundant features. Therefore, the most important sanity check comes from the domain expert who picks the features, models the confounders, and interprets the results.

### Modelling confounders

In order for the algorithm to function properly, all confounding effects must be modelled correctly and completely. Specifically, `sBayes` assumes that—once universal preference and inheritance have been accounted for—the remaining similarity in the data is due to contact. We will briefly discuss the confounders currently considered in the model and give an outlook on future extensions.

Universal preference helps the algorithm to establish a baseline for chance. `sBayes` learns how often a feature is expected in extant languages. There are different conceptual approaches for estimating universal preference, yielding a nuanced interpretation for contact and contact areas. When the baseline is derived from the data alone, it encodes preference in the study area. This is appropriate for a sufficiently large and balanced sample, while small and unbalanced samples are likely to yield a biased baseline, resulting in biased areas. For example, the 30 languages coded for in the Balkans case study are similar precisely because they share a common history, in which case it makes sense to inform the baseline with an empirical prior encoding preference outside the biased sample.

Inheritance helps the algorithm to establish a baseline for chance in a family. There are different conceptual levels (and levels of granularity) at which information about common ancestry can be passed to `sBayes` (Fig. 7). When no information about common ancestry is available, the model does not distinguish between inheritance and contact. Instead, it identifies areas of unspecified shared history, i.e. subsets of languages with similar features whose similarity is only poorly explained by universal preference. When common ancestry is modelled at the family level, `sBayes` estimates one set of probability vectors per language family, picking up contact across families, but not within. When modelled at the clade level, `sBayes` estimates one set of probability vectors per sub-clade of a language family, revealing contact both across families and across clades. It is up to the analyst to define the granularity at which the phylogeny is split into clades: the finer the splits, the more the model is able to pick up contact between closely related languages. However, increasing granularity brings about decreasing statistical robustness. Too few languages per clade (*<* 5) make it difficult to estimate robust probability vectors. In reality, inheritance is a hierarchical process. While all languages in a family are expected to inherit some shared features, close relatives do so more than distant ones. A phylogenetic likelihood could capture this hierarchical process in a principled way. We plan to extend `sBayes` and implement Felsenstein’s pruning algorithm [28], which will compute a tree-based likelihood whenever the user provides a phylogeny for a language family. This will result in better estimates for confounding, making it possible to pick up nuanced signals of contact across and within families.

**Figure 7:**
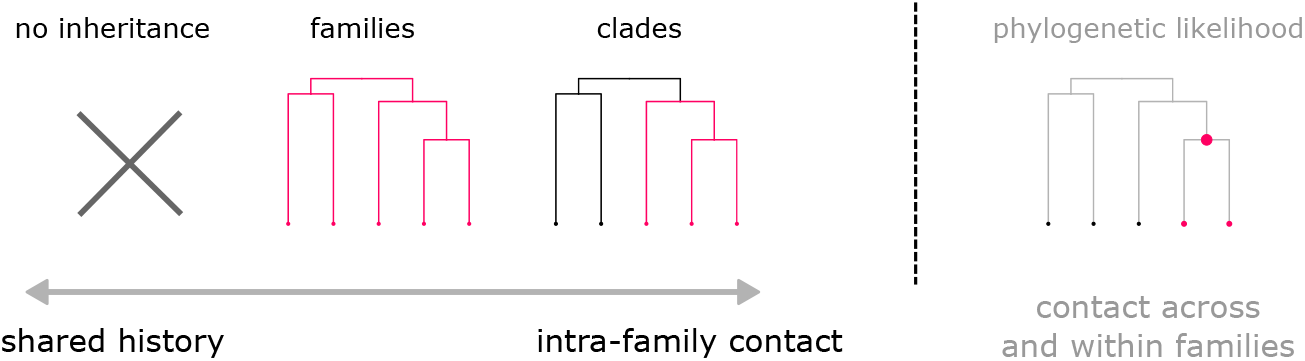
Information about inheritance can be passed to `sBayes` at different levels, causing the algorithm to pick up different contact signals, which range from (unspecified) shared history to intra-family contact. For future versions, a phylogenetic likelihood could model inheritance as a hierarchical process and reveal nuanced traces of contact.

Besides universal preference and inheritance, there are other confounders that could shape the distribution of linguistic features. For example, climate [27], altitude [26], genetics[18], subsistence [6] and population size [35] have all been hypothesised to influence the human sound inventory. To account for the influence of, e.g. climate, we would add an additional effect to the mixture model, assign languages to climate regions and estimate a distribution for each. While adding effects is straightforward, it requires careful consideration. Climate and contact are likely correlated: geographically close languages tend to have a similar climate and they are more likely to be in contact. Thus, a climate confounder would explain parts of the actual contact signal, which might be undesirable.

### Testing hypothesis of spatial evolution

The geo-prior models the prior belief of areas as a function of costs to traverse geographical space: What is the probability that languages have been in contact given the distance between them? There are two different applications of the geo-prior. First, it helps to guide inference. An informed geo-prior will encourage the algorithm to delineate spatially compact areas, coinciding with traditional ideas of what constitutes a linguistic contact area. A reasonably informed geo-prior penalises but does not exclude: if the contact signal is strong enough in remote languages, the algorithm will still report the similarities between them as areas. Second, the geo-prior can be used to test hypotheses of spatial evolution. For instance, in the dense vegetation of the Amazon rain forest contact might be more likely between languages connected by navigable waterways. One could define a model with a uniform geo-prior and one with a strong geo-prior with costs defined as canoeing distance along the river network. The marginal likelihood, e.g. approximated with a stepping stone sampler [73], could quantify the evidence of each model. Bayesian model selection [30] could determine which model is more likely given the data. In a similar way it is possible to model other prior beliefs about geography, socioeconomics, or environment and test their influence on the clustering: are emerging contact areas best explained by hiking effort, trade routes, or vegetation?

### Applications beyond linguistics

Besides language contact there are other domains where `sBayes` can be applied. Contact between groups has many more dimensions than language, which can be analysed using `sBayes` as long as they can be captured in the form of features. One dimension is culture: wherever people are in contact, they tend to exchange artefacts, but also cultural practices, ideas, rituals, mythology, etc. All of these types of exchange may leave traces in the anthropological and archaeological record. Although feature-based interpretations of cultural practices have been criticised [43], there is an ongoing tradition to do so (see e.g. [55, 53, 39]). Studies conducted show that meaningful reconstructive models can be built on the basis of cultural features [61, 49, 39, 63, 70]. Moreover, the geo-prior could be used to test for hypotheses of evolution in space and compare human evolution across different dimensions. Does cultural contact follow similar pathways as genetic variation? This hypothesis could be evaluated against empirical data by using spatial clusters emerging in genetic data as a geo-prior when applying `sBayes` to cultural data. Potentially, the use of `sBayes` might also be explored to tackle other problems outside the broader domain of cultural evolution. In ecology, for example, `sBayes` might reveal ecological habitats while controlling for preferences due to confounders such as climate or soil patterns. In environmental science, `sBayes` might show toxic hotspots while controlling for known effects due to population density or traffic. In social network data the proposed algorithm might reveal similarities across users, while controlling for socio-cultural preferences.

## Auhtors’ contribution

PR, RG and NN conceived the idea, NN and PR developed the methodology, implemented the algorithm and carried out the case studies. BB and RG framed the model in terms of theories of linguistic distribution. RG and PM collected the data for the South American case study and interpreted the results. AE and BS collected the data for the Balkans case study and interpreted the results. PR and NN led the writing of the manuscript. BB, RW, RG, AE and BS contributed critically to the methodology and the draft. All authors gave final approval for publication.

## Data availability

The data for the two case studies are available at https://github.com/derpetermann/sbayes, together with the software, the installation guidelines and a manual.

## Notes

https://github.com/derpetermann/sBayes

### Competing Interest Statement

The authors have declared no competing interest.

