## Supporting Information for "Contact-tracing in cultural evolution: a Bayesian mixture model to detect geographic areas of language contact"

March 31, 2021

#### Contents

**S1 A brief history of contact linguistics** **S2**
S1.1 Qualitative approaches . . . . . S2
S1.2 Quantitative approaches . . . . . S3

**S2 Likelihood** **S3**

**S3 Priors** **S5**
S3.1 Universal preference and preference in a family . . . . . S5
S3.2 Size and number of areas . . . . . S6

**S4 Ranking areas** **S6**

**S5 Sampling** **S6**

**S6 Software** **S8**

**S7 Simulation Study** **S8**

**S8 Case Study: Western South America** **S13**

**S9 Case Study: Balkans** **S20**

This document provides supporting information for the main article "Contact-tracing in language evolution".

### S1 A brief history of contact linguistics

#### S1.1 Qualitative approaches

There are two principled ways to approach the detection and description of a linguistic area: bottom-up and top-down [Muysken, 2008, Muysken et al., 2014]. The starting point for the bottom-up approach is an observation of one or more salient characteristics for a particular geographical area, which is likely due to contact. These characteristics may be linguistic in nature, but also e.g. cultural traits. This may give an impetus to look at the distribution of (further) linguistic features in the same approximate region, until a picture emerges of a (generally fuzzy) area where languages share a set of features. The 'discovery' of the Balkan Sprachbund was the result of such a process [Friedman, 2011]. Ideally (though certainly not always) this is coupled with data from other disciplines, such as archaeology or anthropology that also suggest a history of contact [Bickel and Nichols, 2006, Campbell, 2006, van Gijn, 2020, Bickel, 2020].
The advantage of the bottom-up approach is that the proposal of a linguistic area is usually firmly embedded in solid regional and genealogical expert knowledge. And if carried out properly, a fairly complete picture of historical socio-dynamics can emerge, and because of the intimate knowledge of the researcher with the languages in the area, subtle signals can be picked up. The obvious disadvantage is that the quality of the results depends heavily on the expertise of the researcher, and/or accidental observations. Moreover, this approach is open to criticism of selective attention: by cherry-picking features that are suggestive of diffusion, a false or exaggerated image of significant areal convergence can be conveyed.
The other way to approach linguistic areas, is top-down. Here the approach is to assemble a principled set of features, and screen the distribution of the feature values for areal clustering, either with or without a previous areal hypothesis. This type of approach has generally given rise to more quantitatively oriented approaches (see below). Examples of qualitative top-down approaches are most often found in large-scale surveys of feature distributions. The main technique for discovering areal clusters of features in this approach is eyeballing a feature distribution map. Many contributions to Dryer and Haspelmath [2013] are of this nature, as are e.g. Haspelmath [2001], Heine [2008], Krasnoukhova [2012], Miceli and Dench [2017] on a continental level.
The advantage of the top-down over the bottom-up approach is that it reduces the dependency on areal expertise and concomitant subjectivity somewhat (the *a priori* choice of features remains relatively arbitrary, though). One of the disadvantages is that it may miss contact signals that would have been picked up in a bottom-up approach, and thus it may underestimate the contact signal. Moreover, the exploratory and incomplete nature of this approach is not always recognised as such and thus this approach runs the risk that results are not followed up by more detailed and multi-disciplinary research, and so that it may lose the historical embedding present in bottom-up approaches [Campbell, 2006, Van Gijn and Wahlström, forthcoming].
Either approach needs to deal with a number of highly problematic issues. Van Gijn and Wahlström [forthcoming] list three fundamental problems for determining linguistic contact areas (see also e.g. Masica [2001], Bickel and Nichols [2006], Stolz [2002, 2006], Campbell [2006] for discussion).

1. **The boundary problem**: Establishing the geographical boundaries of a linguistic area is often based on the distribution of features. This is problematic because the distributions of different features rarely overlap completely.

2. **The language problem**: There seems to be no non-arbitrary way to determine the minimum number of languages required to speak of a linguistic area.

3. **The feature problem**: There are no established criteria to determine the diagnostic value of features for particular linguistic areas, nor of the minimum number of features required.

On top of these three basic problems, which relate to the status of linguistic areas as theoretical constructs, there is no principled method of distinguishing between effects of contact and inheritance, so that one is again dependent on highly expert and potentially subjective views. This latter point is particularly problematic for areas with genealogically related languages (see e.g. Dunn et al. [2008], Noonan [2010], Epps et al. [2013], Bowern [2013]), and is the reason behind

the fact that linguistic areas almost always consist of genealogically diverse languages, sometimes even by definition (see e.g. Enfield [2005]). The paradigmatic example of the Balkans is a salient exception, since—apart from Balkan Turkish—all languages belong to the Indo-European family, albeit to several different sub-branches.
The way forward with respect to at least some of these issues is the use of quantitative methods, which have started to surface roughly over the last decade.

##### S1.2 Quantitative approaches

There are two types of quantitative methods that help researchers to analyse language contact in space: methods for hypothesis testing and clustering methods. Hypothesis testing assumes that the analysts have already singled out a potential contact area—based on information from linguistic and non-linguistic disciplines such as geography, archaeology, or genetics—for which they want to assess the robustness and significance of the areal signal. Such an approach was taken in Bickel and Nichols [2006] to test the hypothesis that the Circum-Pacific is a large contact area, extending over both sides of the Pacific Coast. The study finds statistically significant differences between the distribution of features in the contact area, while controlling for both genealogical relatedness and universal preference. A similar study used Bayesian analysis to reveal a gain in linguistic similarity in the British Isles, cross-cutting lines of linguistic ancestry [Dedio et al., 2019].

Statistical testing might address selective attention on the feature level, but not on the language level: a rigorous statistical test might ensure that a signal in a fixed set of objects differs from chance, but it might not itself provide justification for why a specific set of languages was grouped into an area *a priori*, given the myriads of other potential areal groupings in the data. In contrast, clustering methods do not require the analyst to define potential contact areas *a priori*, but offer the promise of delineating them from the data. Clustering is the task of assigning objects to groups, such that the distance (i.e., dissimilarity) within each group is minimised, while the distance between groups is maximised.
Two types of clustering methods have been used in the context of historical linguistics: cost-based clustering and model-based clustering. Cost-based clustering—such as k-means, hierarchical or spectral clustering—minimises a cost function (usually distances) to assign each object to exactly one of $k$ groups. Cost-based clustering was mainly applied to dialect data: k-means for Swedish [Lundberg, 2005], hierarchical clustering for Swiss-German [Scherrer and Stoeckle, 2016], French [Goebl, 2008] and English [Szmrecsanyi, 2011] and spectral clustering for Dutch [Wieling and Nerbonne, 2011]. While these methods address the problem of selective attention, they are not based on probabilistic models and most often discretely assign languages to areas, missing out on the gradual transitions likely to be present in language data [Bickel and Nichols, 2006] and dialect data in particular [Jeszenszky et al., 2018]. Model-based clustering, on the other hand, assumes that each object results from a mixture of two or more unknown distributions, i.e. the clusters. The assignment of objects to groups is probabilistic: an object belongs to several clusters, each with a certain probability. In historical linguistics, many case studies on model-based clustering fall back on the STRUCTURE algorithm, developed to infer population structure from individual genotypes and gene frequencies [Pritchard et al., 2000]: STRUCTURE was applied to cluster Tasmanian [Bowern, 2012], Melanesian [Dunn et al., 2008] and Sahul [Reesink et al., 2009] languages, as well as Finnish [Syrjänen et al., 2016] dialects.
STRUCTURE addresses the issue of gradual transitions, but when applied to non-related languages fails to distinguish among mutually confounding effects: although it can provide important clues for reconstructing the past [Dunn et al., 2008], it essentially cannot distinguish between contact and inheritance Reesink and Dunn [2012], Reesink et al. [2009]. Moreover, STRUCTURE critically builds on the Hardy-Weinberg equilibrium [Hao and Storey, 2019], which has no natural interpretation when applied to languages or cultures.

#### S2 Likelihood

`sBayes` models each feature as coming from a distribution that is a weighted mixture of universal preference, inheritance and contact. The unknown weights — $w_{\text{universal}}$, $w_{\text{inherit}}$ and $w_{\text{contact}}$ — quantify the contribution of each of these three effects. For a single language $l$, a feature $f$ and a state $s$, this yields the following mixture model:

$$
\begin{aligned}
P(X_{l,f} = s|w, \alpha, \beta, \gamma) = w_{\text{universal},f} \cdot P_{\text{universal}}(X_{l,f} = s|\alpha_f) \\
+ w_{\text{inherit},f} \cdot P_{\text{inherit}}(X_{l,f} = s|\beta_f) \\
+ w_{\text{contact},f} \cdot P_{\text{contact}}(X_{l,f} = s|\gamma_f)
\end{aligned} \tag{1}
$$

The **likelihood for universal preference** estimates the probability that the state $s$ of feature $f$ in language $l$ is preferred due to universal preference:

$$P_{\text{universal}}(X_{l,f} = s|\alpha_f) = \alpha_{f,s} \tag{2}$$

In Equation 2, $s$ is modelled as an outcome of a categorical distribution with an unknown universal probability vector $\alpha_f = [\alpha_{f,1}, ..., \alpha_{f,k}]$ where the entries sum to 1. The probability to observe state $s$ in feature $f$ equals $\alpha_{f,s}$. Since the universal distribution is the same globally, the probability vector $\alpha_f$ is shared among all languages. Imagine we aim to find the probability vector that explains the global distribution of palatal nasals as shown in Figure 2b, Main Paper. Palatal nasals are present in four out of sixteen languages. When only considering universal preference, the universal probability vector below would best explain the data:

$$\alpha_{\text{palatal}} = \begin{bmatrix} \alpha_{\text{present}} & = 0.25 \\ \alpha_{\text{absent}} & = 0.75 \end{bmatrix}$$

The **likelihood for inheritance** estimates the probability that language $l$ in family $\varphi$ has inherited the state $s$ of features $f$:

$$P_{\text{inherit}}(X_{l,f} = s|\beta_{f,\varphi(l)}) = \beta_{f,\varphi(l),s} \tag{3}$$

Here, the subscript $\varphi(l)$ denotes the family of language $l$. Again, $s$ is a random outcome of a categorical distribution with an unknown family probability vector $\beta_{f,\varphi(l)} = [\beta_{f,\varphi(l),1}, ..., \beta_{f,\varphi(l),k}]$. In contrast to universal preference, the probability vector depends on the language family: only languages from the same family share a common distribution. `sBayes` estimates one probability vector per family $\varphi$ and evaluates it against all languages in $\varphi$. In other words, $s$ is assumed to be inherited. Imagine we aim to find family probability vectors to explain the distribution of possessive inflection in families $\varphi_{\text{red}}$ and $\varphi_{\text{blue}}$ depicted in Figure 2b, Main Paper. In $\varphi_{\text{red}}$ possessive inflection is obligatory in five out of eight languages, in $\varphi_{\text{blue}}$ in four out of eight. When only considering inheritance, the probability vectors below would best explain the data:

$$\beta_{\text{infl, red}} = \begin{bmatrix} \beta_{\text{obligatory, red}} & = 0.625 \\ \beta_{\text{non-obligatory, red}} & = 0.375 \end{bmatrix} \quad \text{and} \quad \beta_{\text{infl, blue}} = \begin{bmatrix} \beta_{\text{obligatory, blue}} & = 0.5 \\ \beta_{\text{non-obligatory, blue}} & = 0.5 \end{bmatrix}$$

Note that we could also model relatedness on a different than the family level. In particular, we can incorporate closer relatedness by defining the likelihood of inheritance based on clades, in which case the model estimates a separate family probability vector $\beta_{f,\text{clade}(l)}$ for every clade at a certain time depth:

$$P_{\text{inherit}}(X_{l,f} = s|\beta_{f,\text{ clade}(l)}) = \beta_{f,\text{ clade}(l),s} \tag{4}$$

We applied this idea in the case study on the Balkans, where we modelled a separate likelihood for the Slavic, Romance, Albanian, and Greek languages. This is crucial in order to estimate the contribution of contact and inheritance in the evolution of an area of shared features that is characterised by closely related varieties. Spinning this idea further, ideally the confounding effect of inheritance is derived from the entire evolutionary history in a phylogenetic reconstruction. We elaborated on this idea in the Discussion of the Main Paper.

The **likelihood for contact** estimates the probability that the state $s$ of feature $f$ was passed on to language $l$ by contact in area $Z$. `sBayes` proposes a set of contact areas $\mathcal{Z}$ and then evaluates the likelihood for each area $Z \in \mathcal{Z}$:

$$P_{\text{contact}}(X_{l,f} = s|\gamma_{f,Z(l)}) = \gamma_{f,Z(l),s} \tag{5}$$

Here, the subscript $Z(l)$ denotes the contact area of language $l$. The probability vector depends on $Z$: only observations in languages which are assigned to the same area are assumed to originate from a common categorical distribution. `sBayes` estimates one areal probability vector per contact area and evaluates it against all corresponding languages. Each language can only belong to one area at a time, or to no area at all. However, in contrast to inheritance, the assignment of languages to areas is not fixed, but inferred. Returning to the example of numeral base systems, four languages in the area $Z$ have a hybrid base system and one a vigesimal one (Fig.2b, Main Paper). Again, when only considering contact, the areal probability vector below would best explain the distribution in $Z$:

$$\gamma_{\text{base, } Z} = \begin{bmatrix} \gamma_{\text{decimal, } Z} & = 0.0 \\ \gamma_{\text{vigesimal, } Z} & = 0.2 \\ \gamma_{\text{hybrid, } Z} & = 0.8 \end{bmatrix}$$

`sBayes` allows for multiple contact areas $\mathcal{Z} = \{Z_1, Z_2, ...\}$, each with their own set of areal probability vectors.

### S3 Priors

#### S3.1 Universal preference and preference in a family

The more a state is universally preferred (or conversely preferred in a family), the less likely a similar occurrence in $Z$ is regarded as evidence for contact. However, what is rare in our sample (i.e., our study area) might be abundant outside and vice versa. In Fig. S1a most languages in the sample (dashed polygon) have state B (yellow), while outside state A (blue) is preferred. Since the languages in our sample are biased, they lead to a biased estimate for universal preference: $Z$ emerges as a contact area.

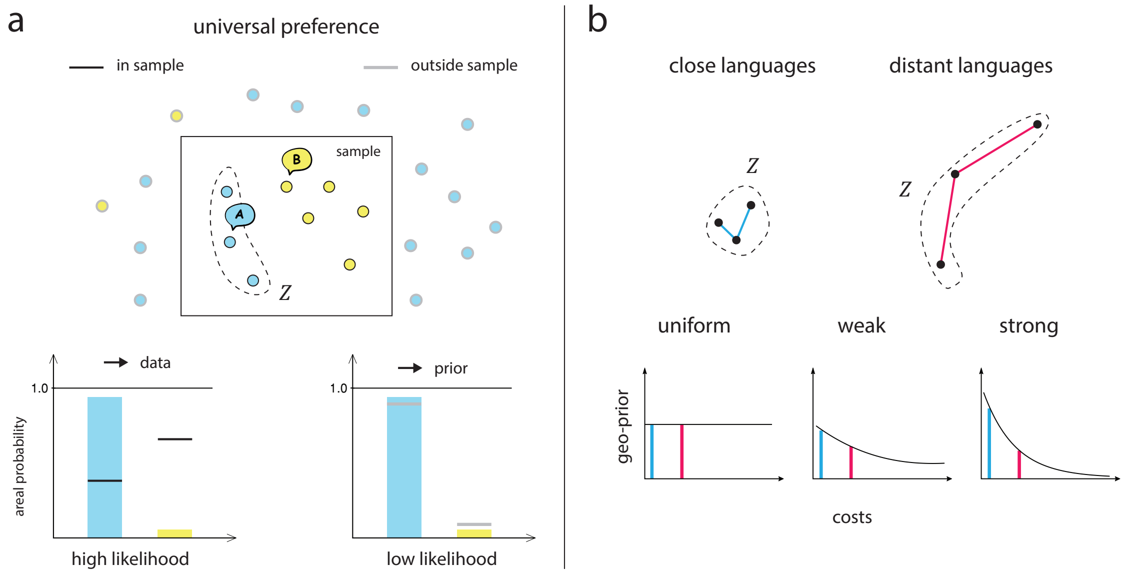


Figure S1: (a) In the sample (i.e., the study area), universal preference is biased towards state $B$. Area $Z$ contains only languages with state $A$. $Z$ has low entropy and differs from universal preference resulting in a high likelihood (left graph). Outside the study area (i.e. in the population of all languages) $A$ is preferred. When used as a prior for universal preference the likelihood decreases, as $Z$ no longer differs from universal preference (right graph). (b) The stronger the geo-prior, the more it favours areas with geographically close languages.

An informed prior allows us to express specific knowledge about universal preference before seeing the data (see equation 8, Main Paper). In the example in Fig S1a, a prior for universal preference is derived from additional data from outside the sample to inform the mean $\mu$ and precision $\rho$ of the prior distribution for universal preference: $\mu_A = 13/15$, $\mu_B = 2/15$ and $\rho = 15$. In this particular case, the prior has the same effect as if all 23 languages were present in the sample (with the important difference that languages outside the dashed polygon cannot be part of an area). Alternatively, $\mu$ and $\rho$ can be estimated from previous experiments or they can be informed from the literature.

##### S3.2 Size and number of areas

In addition to the geo-prior, there are two implicit parameters relating to the prior probability of contact areas:

- $m$ is the size of a single area, that is, the number of languages in $Z$. `sBayes` employs two types of priors for $m$. The *uniform area prior* assumes that each area $Z$ is equally likely a-priori. This puts an implicit prior on size, such that larger $m$ are preferred over smaller ones: there are $(M-m+1)/m$ more ways to choose an area of size $m$ from a population $M$ than one of size $m-1$, assuming that $m \leq M/2$. The *uniform size prior* assumes that all $m$ are equally likely a-priori, i.e. $P(m)$ has a uniform prior in the interval $[\min(m), \max(M)]$.

  For a strong areal signal, the influence of each prior is negligible. For a weak signal, the uniform area prior is more likely to pick up contact traces across many languages, but also runs the risk of overfitting to random noise. We suggest to run the model with both priors and compare model fit (i.e. likelihood) and convergence.

- $n$ is the number of contact areas $Z$ in $\mathcal{Z}$. There is no prior for $n$. Instead, we run the model iteratively and increase the number of areas per run. Then, we compare the performance across $n$ in post-processing.

#### S4 Ranking areas

In post-processing, `sBayes` ranks all areas $n$ according to their posterior evidence for contact. Evidence for contact is expressed as the relative posterior probability of each area $Z_i$ as compared to that of the remaining areas $Z_j$, where $j \neq i$. For example, for $n = 4$, the relative posterior probability of area $Z_1$ is evaluated as follows: for each sample in the posterior distribution only area $Z_1$ remains active, while all other areas $Z_2$, $Z_3$ and $Z_4$ are removed, such that the languages in these areas are only explained by universal preference and inheritance. The mean posterior across all samples with only $Z_1$ active is then compared against the mean posterior with only $Z_2$, $Z_3$, or $Z_4$ active.

#### S5 Sampling

This section explains how `sBayes` samples from the posterior distribution $P(\Theta|D)$ to identify potential contact areas. `sBayes` employs a Markov chain Monte Carlo (MCMC) sampler with two types of proposal distributions: a Dirichlet proposal distribution for weights and probability vectors and a discrete, spatially informed proposal distribution for areas.
The weights and probability vectors are points on a probability simplex: they are bounded between $[0, 1]$ and add to 1. This motivates using a **Dirichlet proposal distribution**:

$$q(\theta_{\text{new}}|\theta_{\text{old}}) = \text{Dir}(\psi_i = 1 + \theta_{\text{old},i} \cdot \kappa_i) \quad \text{for } i \in 1, \dots k \tag{6}$$

The probability simplex has highest density at the current sample $\theta_{\text{old}}$. The pseudo-counts $\kappa$ control the width of the proposal distribution: how far from the current sample should new candidates $\theta_{\text{new}}$ be recruited. Candidates are accepted with Metropolis-Hastings probability:

$$\min(1, \frac{P(\theta_{\text{new}})}{P(\theta_{\text{old}})} \cdot \frac{\mathcal{L}(\theta_{\text{new}})}{\mathcal{L}(\theta_{\text{old}})} \cdot \frac{q(\theta_{\text{new}}|\theta_{\text{old}})}{q(\theta_{\text{old}}|\theta_{\text{new}})}), \tag{7}$$

where $q(\theta_{\text{old}}|\theta_{\text{new}})$ is the back probability to move from $\theta_{\text{new}}$ back to $\theta_{\text{old}}$:

$$q(\theta_{\text{old}}|\theta_{\text{new}}) = \text{Dir}(\psi_i = 1 + \theta_{\text{new},i} \cdot \kappa_i) \quad \text{for } i \in 1, \dots k \tag{8}$$

The Dirichlet distribution is not symmetric and proposal and back probability are not generally equal.

The assignment of languages to areas $Z$ is discrete, that is, languages belong to a contact area or do not. Hence, continuous proposal distributions are not applicable. Moreover, the number of possible contact areas grows exponentially with the number of languages in the sample: for 300 languages there are more ways to randomly assign points to $Z$ than atoms in the universe. At the same time we would not expect language contact to be purely random either, but spatially clustered. Thus, a purely random proposal distribution is neither very efficient, nor very smart.

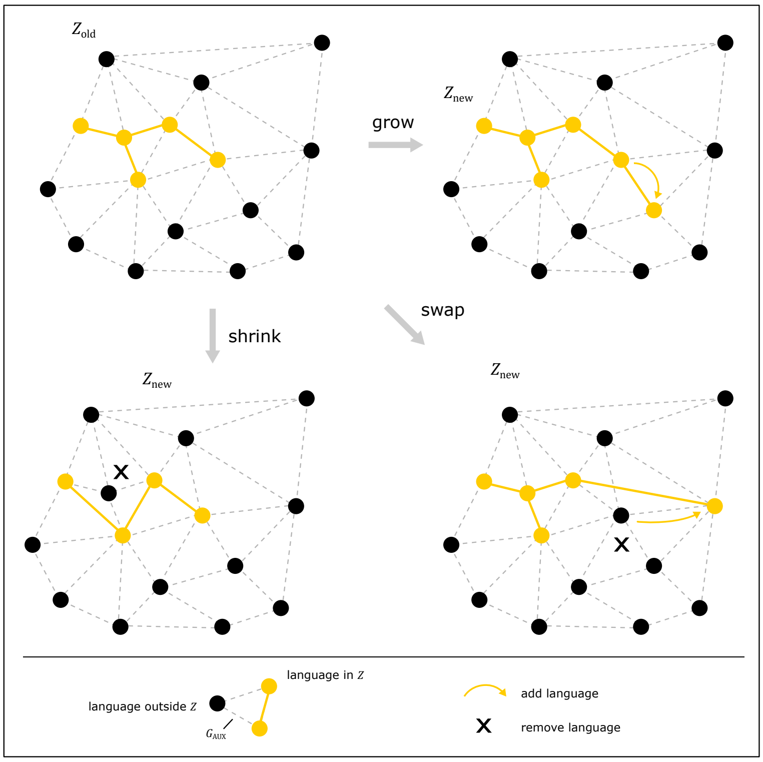


Figure S2: The spatially-informed proposal distribution grows, shrinks or swaps an initial area $Z_{\text{old}}$. In the example, all steps are restricted to the adjacent neighborhood of the initial area defined by the neighborhood graph $G_{\text{AUX}}$.

Instead, `sBayes` employs **a spatially-informed proposal distribution**, with both purely random steps and steps restricted to the adjacent neighbourhood of the current area. The proposal distribution builds on the assumption that languages in $Z$ are likely spatial neighbours, but is flexible enough to relax this assumption when necessary.

We connect the spatial locations of all languages with a Delaunay triangulation, creating an auxiliary neighbourhood graph, $G_{\text{AUX}}$, which captures spatial adjacency. First, a random set of adjacent languages are assigned to an initial contact area $Z_{\text{old}}$. Then, the MCMC repeatedly takes one of the following steps to propose candidates $Z_{\text{new}}$ (Fig. S2):

- the *grow* step adds a language $l$ to $Z_{\text{old}}$.
- the *shrink* step removes a language from $Z_{\text{old}}$. Note that the new candidate area might or might not be a subgraph of $G_{\text{AUX}}$. The Delaunay triangulation facilitates the proposal of contact areas, but does not restrict it.
- the *swap* step first shrinks and then grows $Z_{\text{old}}$.

Growing, shrinking and swapping, the MCMC meanders through the universe of possible contact areas. Some of the steps are to adjacent languages only, and thus spatially-informed by $G_{\text{AUX}}$, others are purely random. The MCMC accepts new candidate areas with Metropolis-Hastings probability:

$$\min(1, \frac{P(Z_{\text{new}})}{P(Z_{\text{old}})} \cdot \frac{\mathcal{L}(\mathcal{Z}_{\text{new}})}{\mathcal{L}(\mathcal{Z}_{\text{old}})} \cdot \frac{q(Z_{\text{new}}|Z_{\text{old}})}{q(Z_{\text{old}}|Z_{\text{new}})}), \tag{9}$$

The ratio of proposal and back probability depends on whether $Z_{\text{old}}$ is grown or shrunk and whether the step is random or to adjacent languages (see Table S1).

The proposal probability for growing depends on how many languages an area can grow to. If growth is restricted to adjacent languages, it is the inverse of $|\text{neighbors}(Z_{\text{old}})|$, the number of neighbours; for the random step the inverse of $\neg Z_{\text{old}}$, the number of languages currently not assigned to any area. The proposal probability for shrinking depends on how many languages an area can drop. It is the inverse of the current size, $|Z_{\text{old}}|$. The back probability relates to the same operations, but reversed: what has grown is shrunk, what has shrunk is grown and $Z_{\text{old}}$ and $Z_{\text{new}}$ switch position. A shrink step is always rejected when the language removed from the

| | *grow* $Z_{\text{old}}$ | | *shrink* $Z_{\text{old}}$ | |
|---|---|---|---|---|
| | random | adjacent | random | adjacent |
| $\frac{q(Z_{\text{new}}\|Z_{\text{old}})}{q(Z_{\text{old}}\|Z_{\text{new}})} =$ | $\frac{\lvert Z_{\text{new}}\rvert}{\lvert\neg Z_{\text{old}}\rvert}$ | $\frac{\lvert Z_{\text{new}}\rvert}{\lvert\text{neighbors}(Z_{\text{old}})\rvert}$ | $\frac{\lvert\neg Z_{\text{new}}\rvert}{\lvert Z_{\text{old}}\rvert}$ | $\frac{\lvert\text{neighbors}(Z_{\text{new}})\rvert}{\lvert Z_{\text{old}}\rvert}$ |

Table S1: Proposal and back probability of the spatially-informed proposal distribution.

area remains without a neighbour in $Z_{\text{old}}$, since reversing this step is not possible. For swapping, the ratio of proposal probability and back probability is 1 if shrinking is followed by growing.

The approach outlined above is tailored to sample from possibly complex, multi-modal discrete distributions with spatial autocorrelation. Shrinking, growing, and swapping explore the auxiliary graph locally, while random steps enable global switches to different macro-regions.
In warm-up, multiple independent chains $C_1, C_2, ...$ explore the parameter space in parallel and find regions of high density from where the main sampler starts. This reduces the risk of the main sampler getting stuck in a local optimum. We plan to follow through on this idea and implement Metropolis Coupled Markov Chain Monte Carlo ($MC^3$) sampling Altekar et al. [2004] in the future.

### S6 Software

`sBayes` is available as a module for Python 3 and a command line tool. The complete source code can be found online at `https://github.com/derpetermann/sbayes`, together with installation guidelines, a manual and the case studies.

A typical analysis in `sBayes` consists of five steps, which are explained in more detail in the manual:

1. Data coding and preparing the features file
2. Defining the priors
3. Setting up and running the MCMC
4. Summarising and visualising the posterior sample

sbayes relies on `Tracer` [Rambaut et al., 2018] to assess the effective sample size (ESS), that is the number of effectively independent draws from the posterior distribution, and the convergence of the sampler. We found that for most experiments, roughly 3 million iterations and 10,000 samples resulted in an ESS of well-above 200. The ESS depends on the number of parameters, i.e. the number of features, language families and contact areas. More complex models with more parameters might require considerably longer sampling.
In addition, `sBayes` can also be used for simulation of linguistic areas and linguistic families.

### S7 Simulation Study

The complete source code of the simulation study can be found on GitHub ( `https://github.com/derpetermann/sbayes`.)

***Experiment 1*** shows that `sBayes` correctly identifies contact in simulated data (Fig. S3). We manually assigned a subset of languages to a single area and simulated contact by aligning some of the features, thus increasing the commonalities between the languages in the area. We varied both the shape and size of the area and the intensity of contact. High intensity corresponds to strong similarities for many features, low intensity to weak similarities for few features. Figure S3 shows the results of one experiment, where the area had the shape of a flamingo and contact was simulated with medium intensity. The posterior distribution of the inferred area spatially overlaps with the simulated one (a). The MCMC converges and the log-likelihood of the inferred area approaches that of the simulated one (b), both recall and precision approach 100% (c).

When the intensity of simulated contact is low, the algorithm is likely to pick up stronger random signals elsewhere in the data: in a set of objects with randomly assigned features, some will necessarily be similar. Whenever this happened, the likelihood of the random signal was always higher than that of the simulated area.

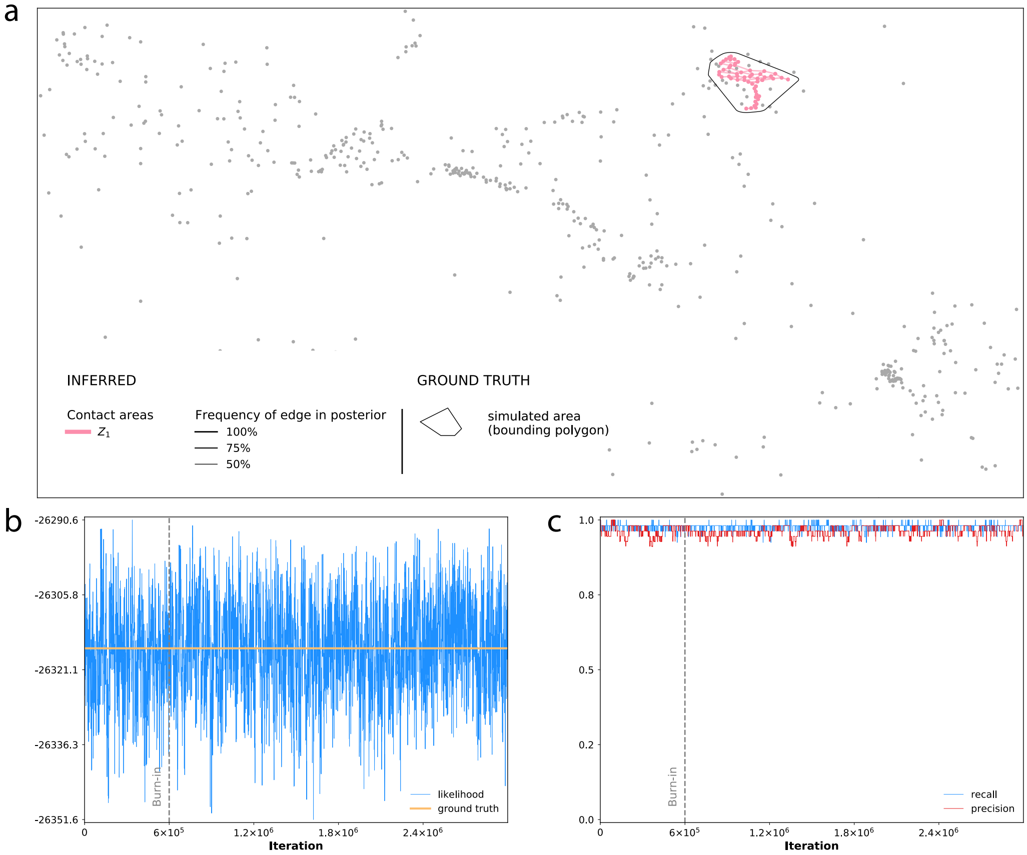


Figure S3: `sBayes` infers a single contact area in simulated data. (a) The posterior distribution of $Z_1$ (pink dots and lines) spatially overlaps with the simulated area (black bounding polygon). The grey dots are the spatial locations of all simulated languages. Languages in $Z_1$ are connected with a Gabriel graph. Line thickness corresponds to the frequency of an edge in the posterior. (b) The trace of the log-likelihood converges and matches the true log-likelihood of the simulation. (c) Both precision and recall approach 100%: `sBayes` has correctly identified the simulated contact area.

A detailed description of ***Experiment 2*** can be found in the main article.

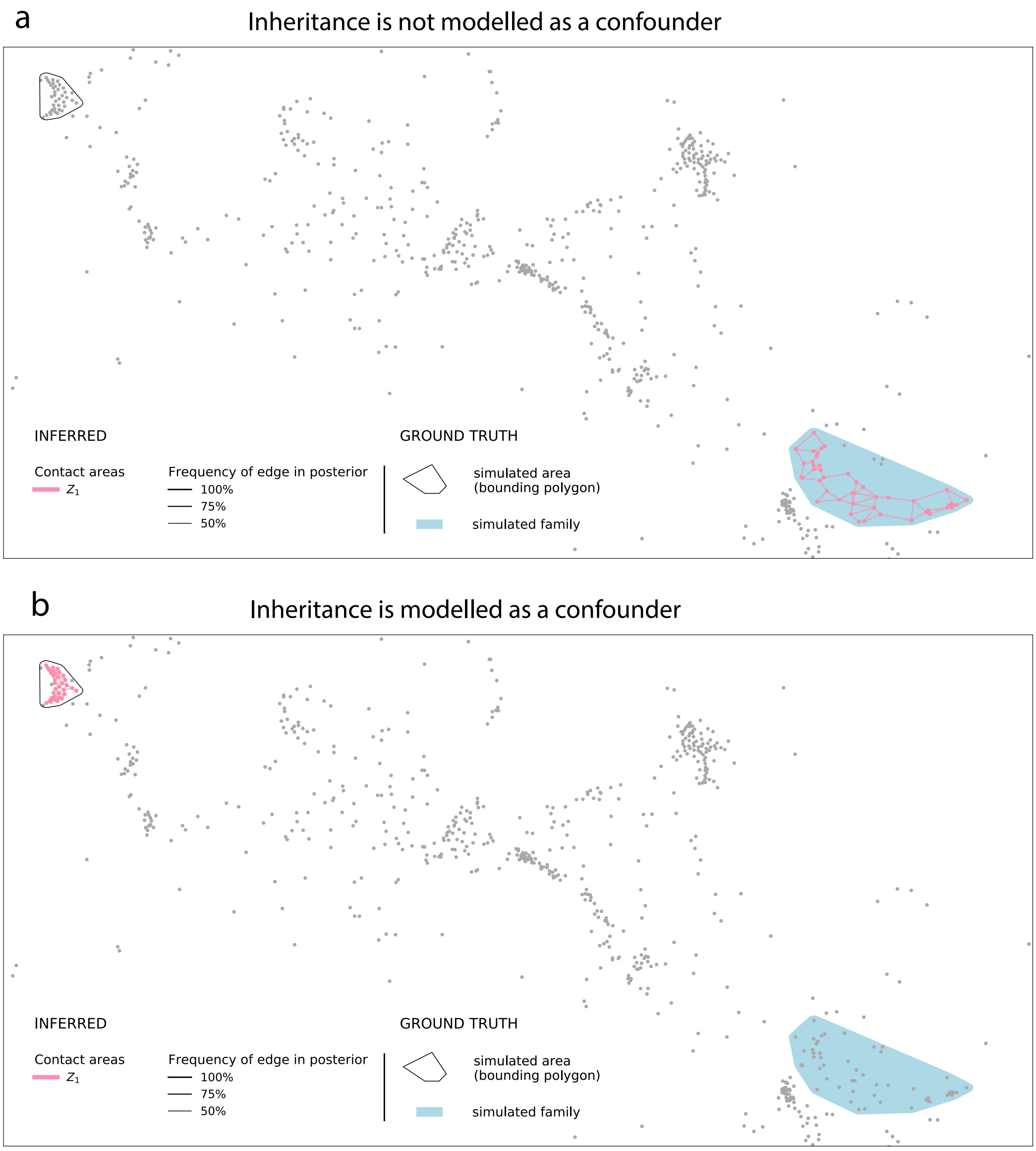


Figure S4: A simulated language family (light blue polygon) and a contact area (black bounding polygon to the left). The grey dots are the spatial locations of all simulated languages. Languages in $Z_1$ are connected with a Gabriel graph, line thickness corresponds to the frequency of an edge in the posterior. (a) When inheritance is not modelled as a confounder, the similarity between languages in the light blue polygon is falsely attributed to contact. For $n = 1$ the posterior of $Z_1$ spatially overlaps with the simulated language family. (b) When inheritance is correctly modelled as a confounder, `sBayes` learns that the similarities in the light blue polygon are best explained by shared descent. The posterior correctly returns the simulated contact area.

***Experiment 3*** illustrates that `sBayes` correctly estimates the number of areas in simulated data. We simulated four contact areas with medium intensity (Fig. S5). We ran `sBayes` and increased the number of areas $n$ with each run. The DIC levels off for $n = 4$, correctly reporting four contact areas (b). For $n = 4$, the areas returned by the posterior perfectly overlap with the simulated areas (a), and both recall and precision reach 100% (c).

***Experiment 4*** demonstrates how an empirically informed prior on the probability vectors for universal preference and inheritance – $\alpha_f$ and $\beta_f$ – allows `sBayes` to robustly identify contact, even in the presence of only few languages. We simulated a contact area in the entire data set. We then ran `sBayes` on a small subset of the data (dashed rectangle in Fig. S6), simulating a situation where only limited and potentially biased data are available for analysis. Again, we tested two setups. For the setup *uniform prior* the prior is flat and all probability vectors are

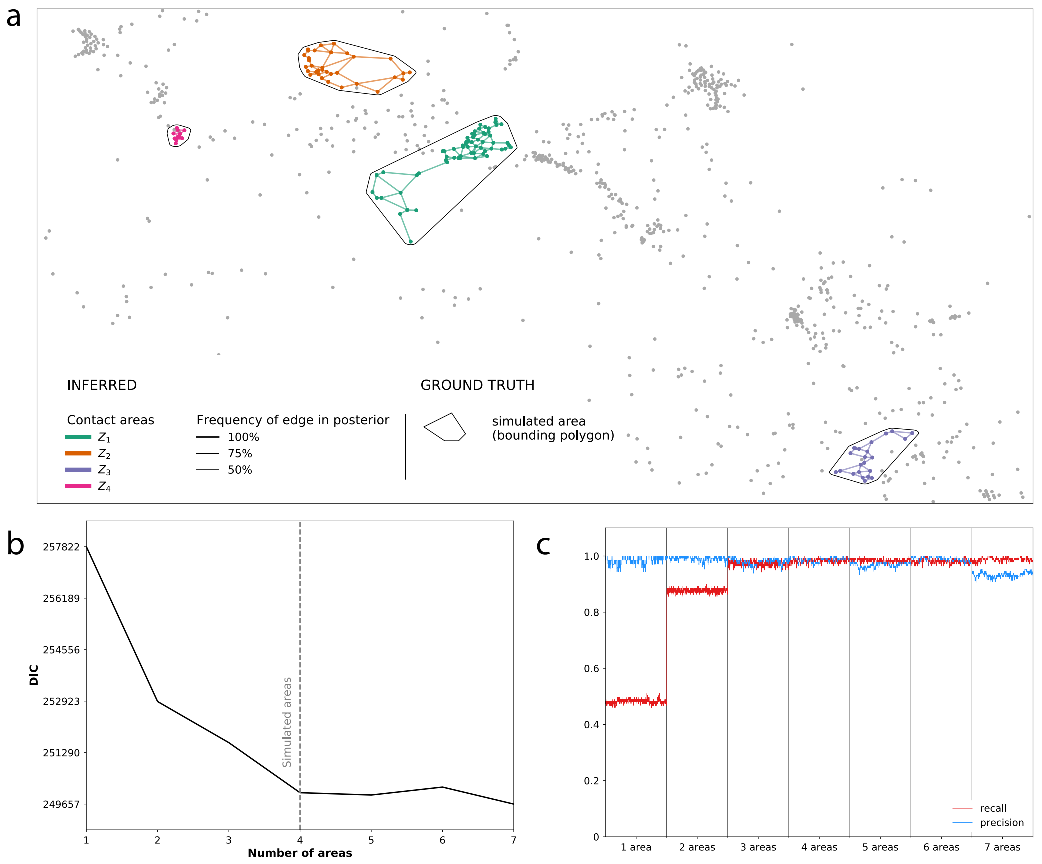


Figure S5: Several contact areas. (a) The posterior distribution consists of areas $Z_1$, $Z_2$, $Z_3$ and $Z_4$ (green, orange, purple and pink dots and lines), which spatially overlap with the four simulated areas (black bounding polygons). The grey dots are the spatial locations of all simulated languages. Languages in each area are connected with a Gabriel graph, line thickness corresponds to the frequency of an edge in the posterior. (b) The DIC levels off for $n = 4$, correctly reporting four areas in the data. (c) For $n = 4$ both recall and precision approach 100%: `sBayes` correctly identifies all simulated areas.

estimated from the small sample alone. `sBayes` fails do report the simulated area (Fig. S6a). The strong signal present in the area is mistaken for universal preference. For the setup *informed prior*, the prior probabilities are informed by all languages initially removed from the sample. `sBayes` correctly returns the simulated area (b).

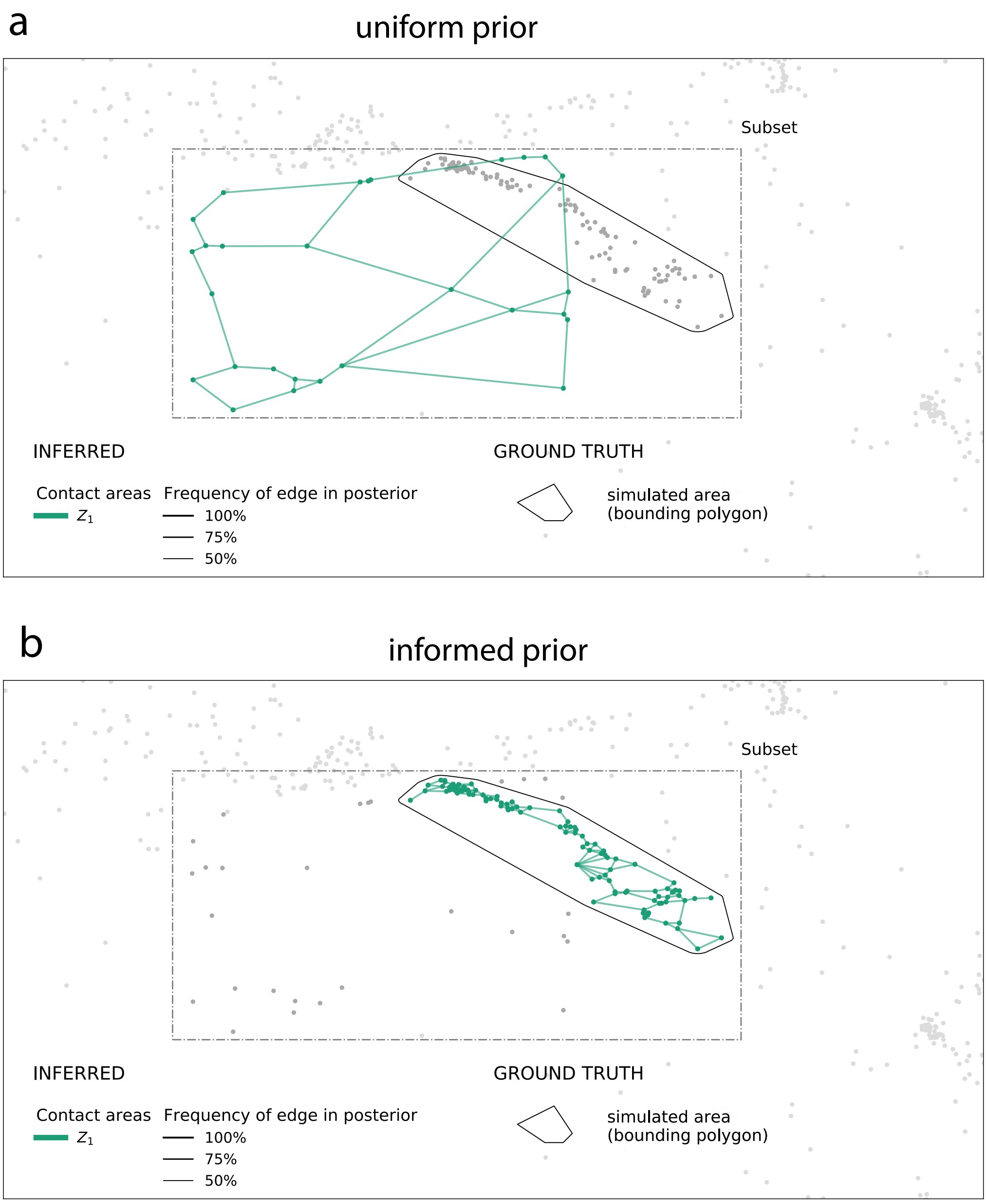


Figure S6: A prior on probability vectors. (a) The grey dots are the spatial locations of all simulated languages. Languages in $Z_1$ are connected with a Gabriel graph, line thickness corresponds to the frequency of an edge in the posterior. The dashed polygon defines the subset passed as input to the algorithm. For the setup *uniform prior* all priors are flat and the posterior distribution of $Z$ fails to report the true contact area. (b) For the setup *informed prior* the prior is empirically informed by all languages outside the sample. The algorithm correctly reports the simulated contact area.

### S8 Case Study: Western South America

For the case study in Western South America, we collected 36 features for a sample of 100 languages in the western Amazon basin and adjacent areas. The features were taken from the literature on the large-scale Andean and Amazonian divide (see van Gijn [2014] for a discussion), as well as literature on smaller linguistic areas in western South America (see Epps and Michael [2017] for an overview). The rationale behind this decision is that we wanted to focus on the distribution of contact signals within the sample of western South American languages.
Figure S7 shows the spatial locations of all languages, Table S2 lists the features used for the analysis. Figure S8 plots the DIC for models with an increasing number of areas $n$.

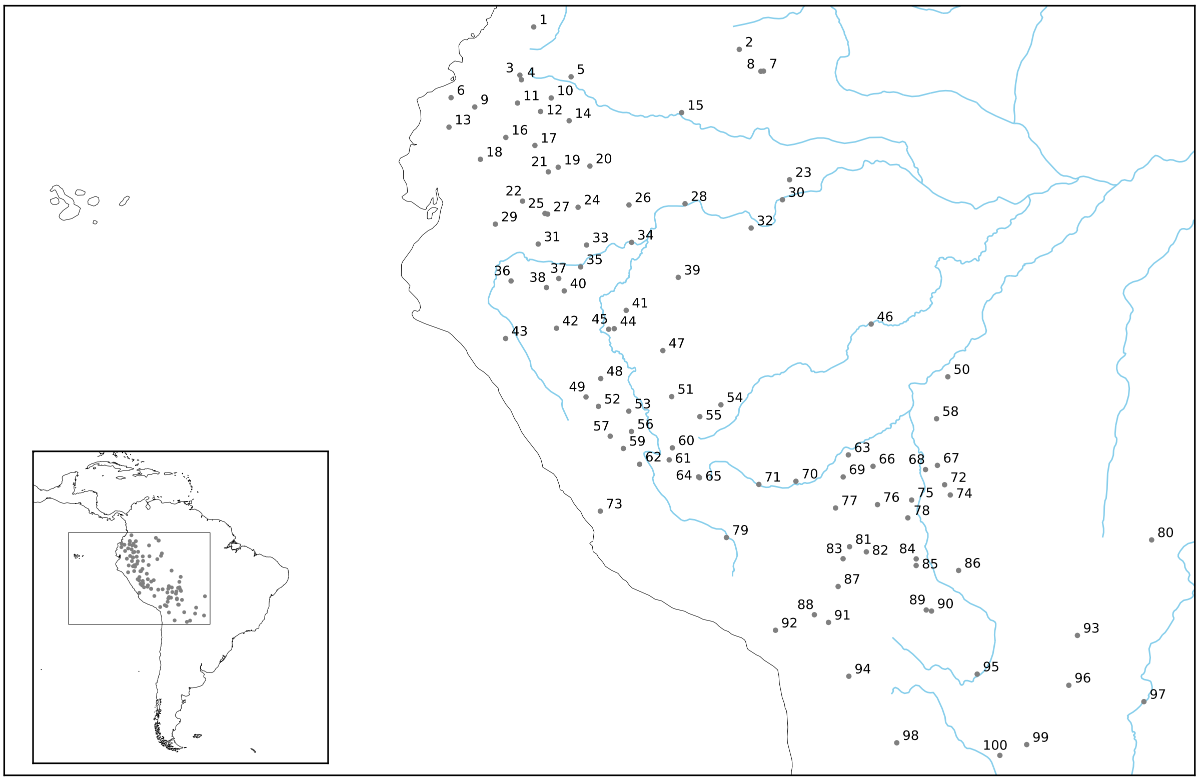

| | | | | | | | | | |
|---|---|---|---|---|---|---|---|---|---|
| 1 | Paez | 21 | Zaparo | 41 | Capanahua | 61 | Caquinte | 81 | Moseten |
| 2 | Cubeo | 22 | Achuar | 42 | San Martin Q | 62 | Nomatsiguenga | 82 | Reyesano |
| 3 | Kamsá | 23 | Miraña | 43 | Cajamarca Q | 63 | Esseeja | 83 | Leko |
| 4 | Inga | 24 | Taushiro | 44 | Shipibo | 64 | Nanti | 84 | Ignaciano |
| 5 | Koreguaje | 25 | Andoa | 45 | Panobo | 65 | Matchiguenga | 85 | Trinitario |
| 6 | Cha'paalachi | 26 | Iquito | 46 | Jarawara | 66 | Chacobo | 86 | Sirionó |
| 7 | Desano | 27 | S Pastaza Q | 47 | Shanenawa | 67 | Kanoe | 87 | Callawaya |
| 8 | Tucano | 28 | Yagua | 48 | Cashibo | 68 | Itene | 88 | Uru |
| 9 | Imbabura Q | 29 | Shuar | 49 | Cholon | 69 | Araona | 89 | Yurakare |
| 10 | Siona | 30 | Omagua | 50 | Karitiana | 70 | Iñapari | 90 | Yuki |
| 11 | Cofan | 31 | Candoshi Shapra | 51 | Ucuyali-Yurua Ash | 71 | Amarakaeri | 91 | Aymara |
| 12 | Napo Quechua | 32 | Matís | 52 | Huallaga Q | 72 | Itonama | 92 | Muylaq Aym |
| 13 | Tsafiki | 33 | Urarina | 53 | Ajy Apurucayali | 73 | Jaqaru | 93 | Chiquitano |
| 14 | Secoya | 34 | Kokama | 54 | Yaminahua | 74 | Baure | 94 | Chipaya |
| 15 | Andoque | 35 | Chamicuro | 55 | Amahuaca | 75 | Cayubaba | 95 | Canichana |
| 16 | Tena Quechua | 36 | Aguaruna | 56 | Pichis Ash | 76 | Cavineña | 96 | Ayoreo |
| 17 | Waorani | 37 | Jebero | 57 | Yanesha' | 77 | Tacana | 97 | Kadiweu |
| 18 | Salasaca Quechua | 38 | Chayahuita | 58 | Wari' | 78 | Movima | 98 | Bol Quechua |
| 19 | N Pastaza Q | 39 | Matsés | 59 | Asheninka | 79 | Cuzco Q | 99 | Chiriguano |
| 20 | Arabela | 40 | Muniche | 60 | Yine | 80 | Paresí | 100 | Tapiete |

Figure S7: All languages coded for the Western South American case study.

Table S2: Features coded in the Western South American case study

| Feature | Description | States |
|---|---|---|
| F1 | Phonemic velar and uvular stops | present, absent |
| F2 | Phonemic /kw/ | present, absent |
| F3 | Phonemic glottalized stops/ejectives | present, absent |
| F4 | Phonemic aspirated stops | present, absent |
| F5 | Phonemic retroflex affricates | present, absent |
| F6 | More phonemic affricates than fricatives | present, absent |
| F7 | Phonemic (bi)labial fricative | present, absent |
| F8 | Phonemic voice contrast for fricatives | present, absent |
| F9 | Phonemic palatal nasal | present, absent |
| F10 | Maximally 1 liquid phoneme | present, absent |
| F11 | Phonemic high central vowel(s) | present, absent |
| F12 | Phonemic front mid versus high contrast | present, absent |
| F13 | Phonemic back mid versus high contrast | present, absent |
| F14 | Phonemic oral-nasal contrast for vowels | present, absent |
| F15 | Morphophonemic nasal spread | present, absent |
| F16 | Contrastive tone | present, absent |
| F17 | Closed syllables: more than 1/3 of consonants can be in coda | present, absent |
| F18 | Distinct ideophone wordclass | present, absent |
| F19 | Clusivity distinction in the pronominal system | present, absent |
| F20 | Noun class/gender distinctions in pronominal system | present, absent |
| F21 | Shape/form-based classifiers | present, absent |
| F22 | Person affixes for possession | present, absent |
| F23 | Morphologically marked alienability distinction in possession | present, absent |
| F24 | Genitive case marking | present, absent |
| F25 | Indigenous monomorphemic numerals above 9 | present, absent |
| F26 | Small (less than 4) case marking system | present, absent |
| F27 | Case marking of A and P roles in transitive clauses | A case, P case, both A and P case, neither A nor P case |
| F28 | Order of adjective and noun | NA, AN, both NA and AN, no adjective class |
| F29 | Marked evidential distinctions on the verb | present, absent |
| F30 | Person prefixes or proclitics | present, absent |
| F31 | Participant roles marked on verb | A marking, P marking both A and P marking, either A or P marking, neither A nor P marking |
| F32 | Morphologically marked isomorphism between intransitive and transitive arguments | S and A, S and P, both S-A and S-P, no isomorphism |
| F33 | Morphologically marked isomorphism of possessor and core verbal argument person markers | present, absent |
| F34 | Constituent ordering of subject and object NPs | OS, SO, both OS and SO |
| F35 | Position verb in transitive sentences | initial, medial, final, free |
| F36 | Switch reference in complex clauses | present, absent |

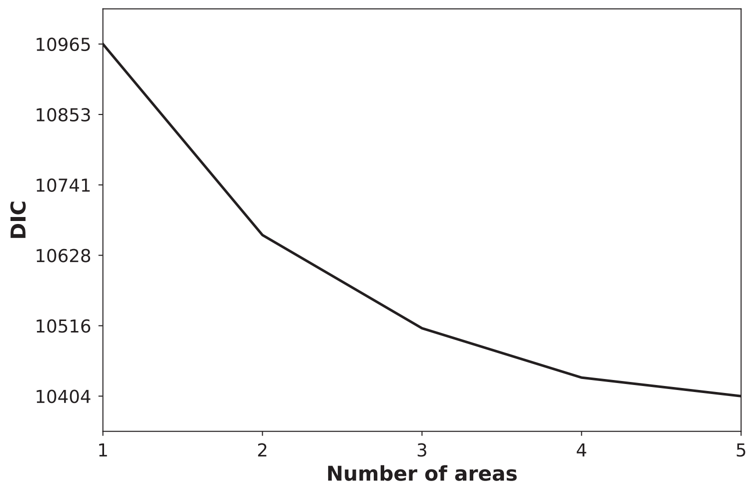


Figure S8: The DIC levels off at $n = 3$, reporting 3 salient contact areas in the Western South American Case study.

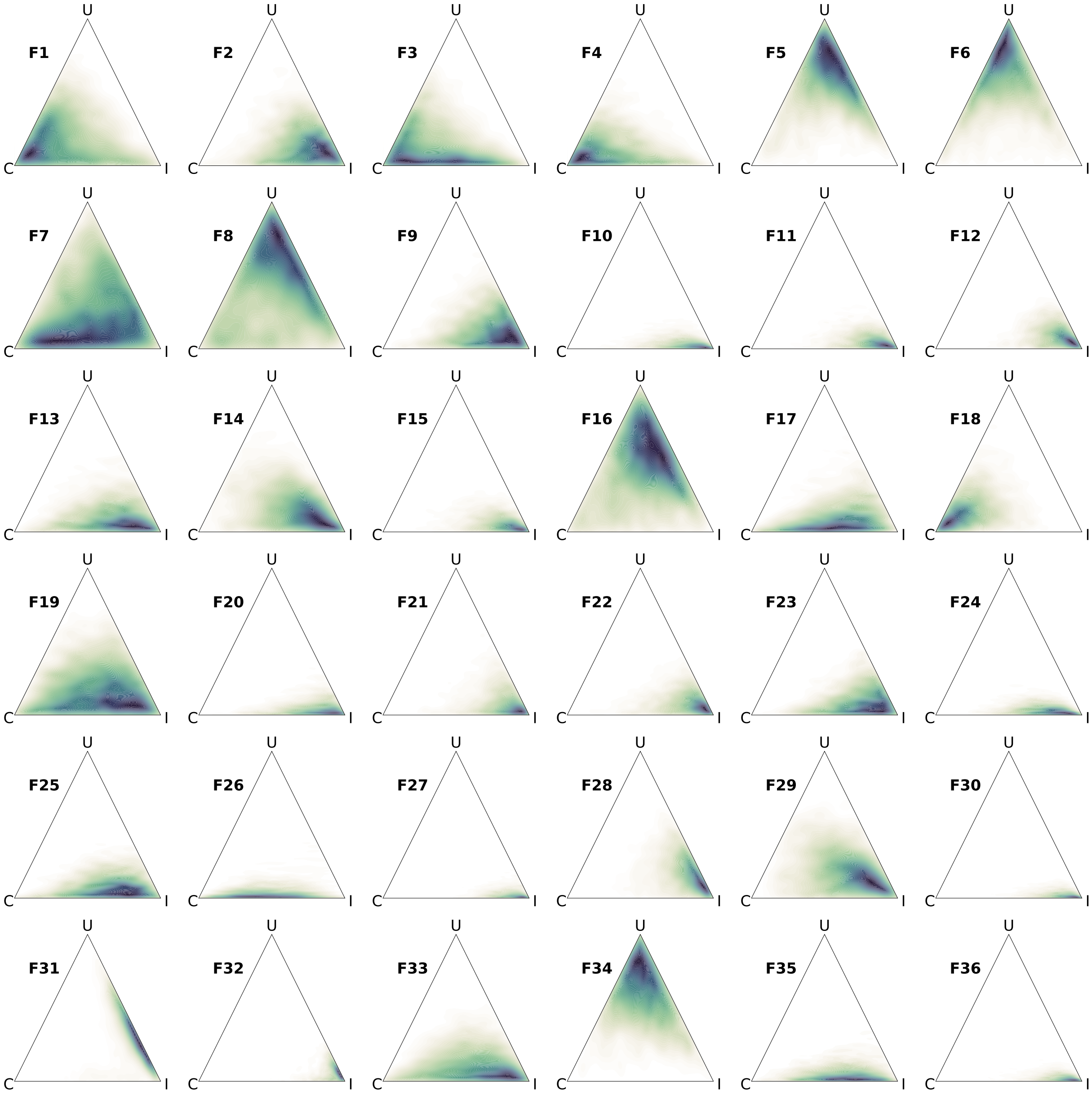


Figure S9: Western South America: weights for all features.

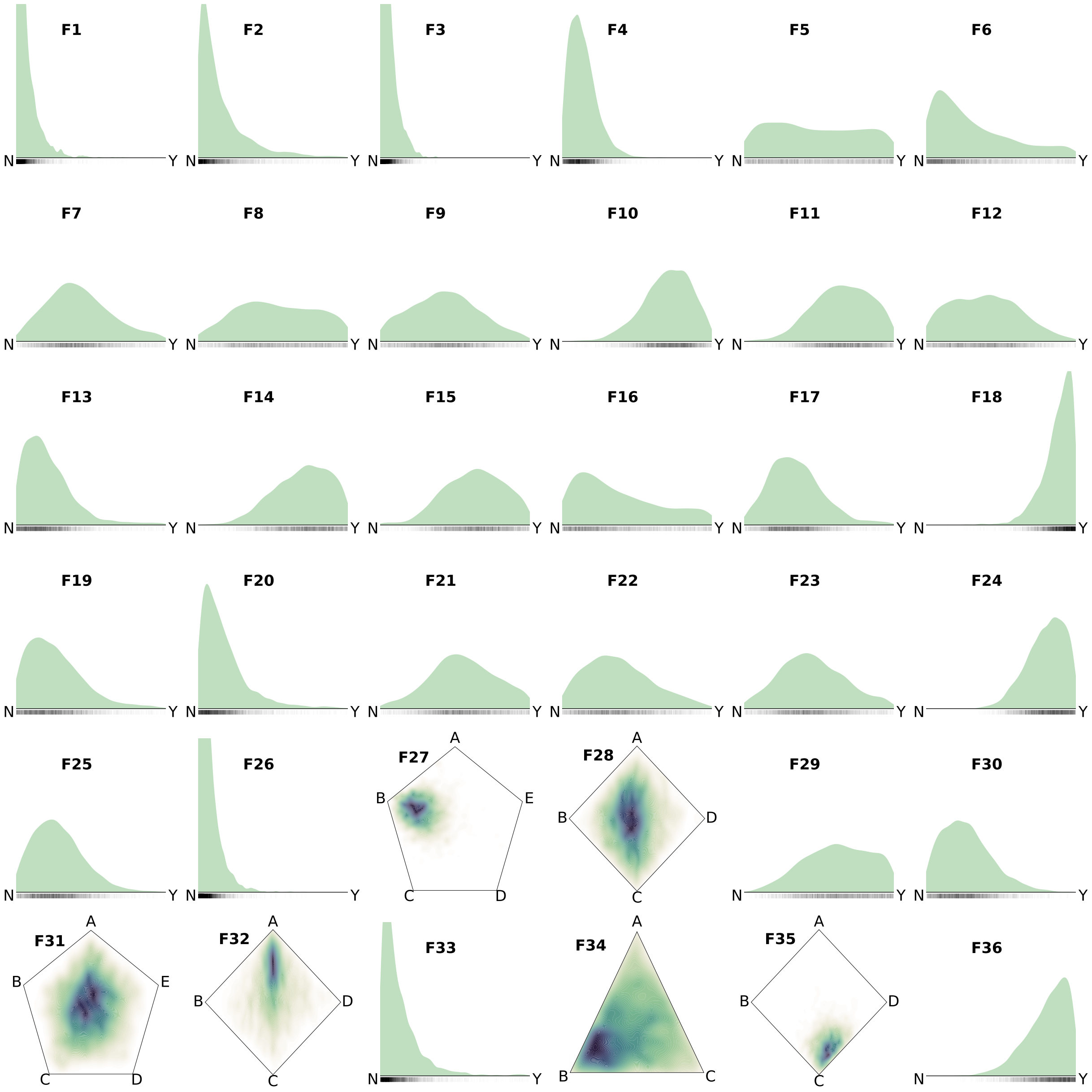


Figure S10: Western South America: areal probabilities in $Z_1$ (the green contact area in Fig. 5a, Main Paper).

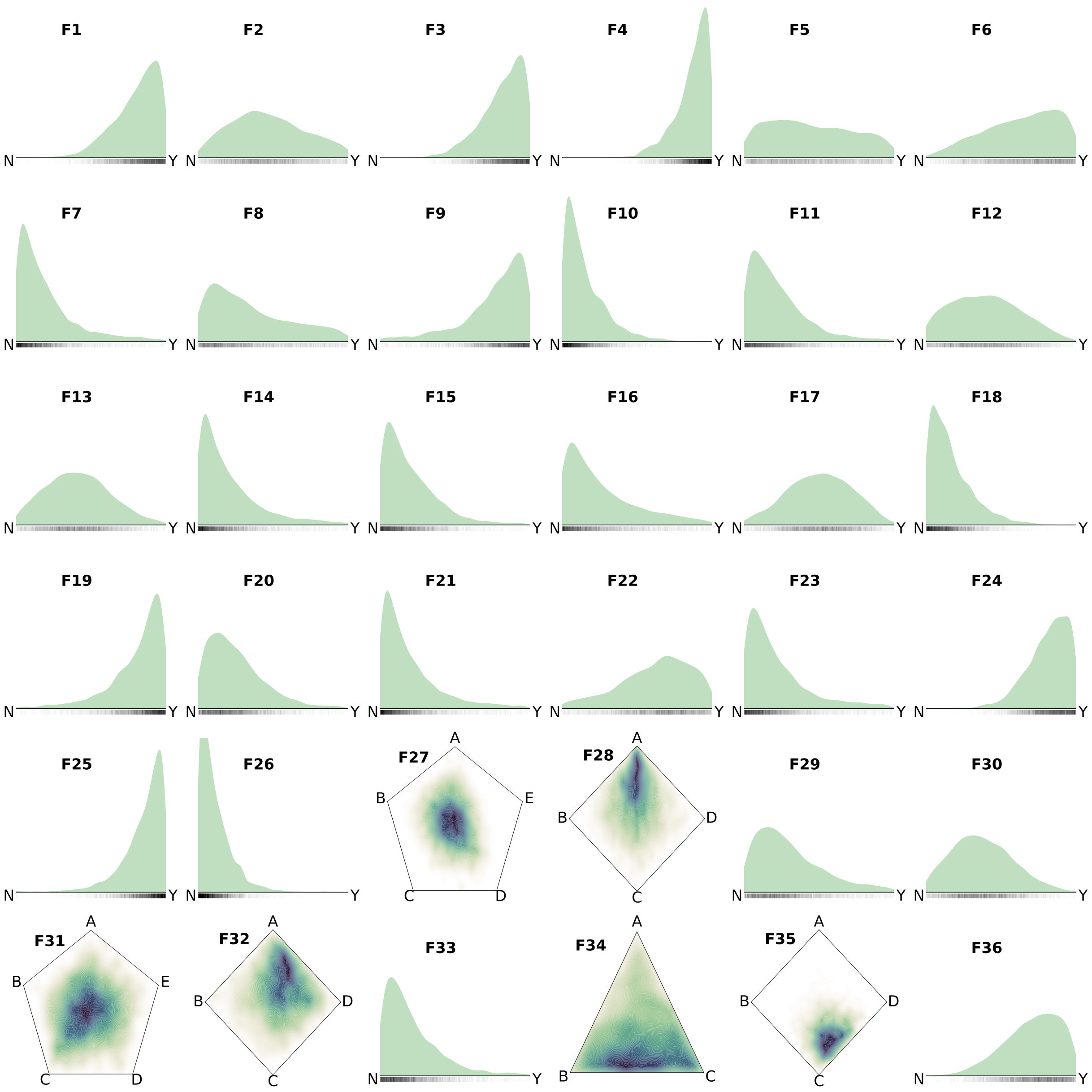


Figure S11: Western South America: areal probabilities in $Z_2$ (the orange contact area in Fig. 5a, Main Paper).

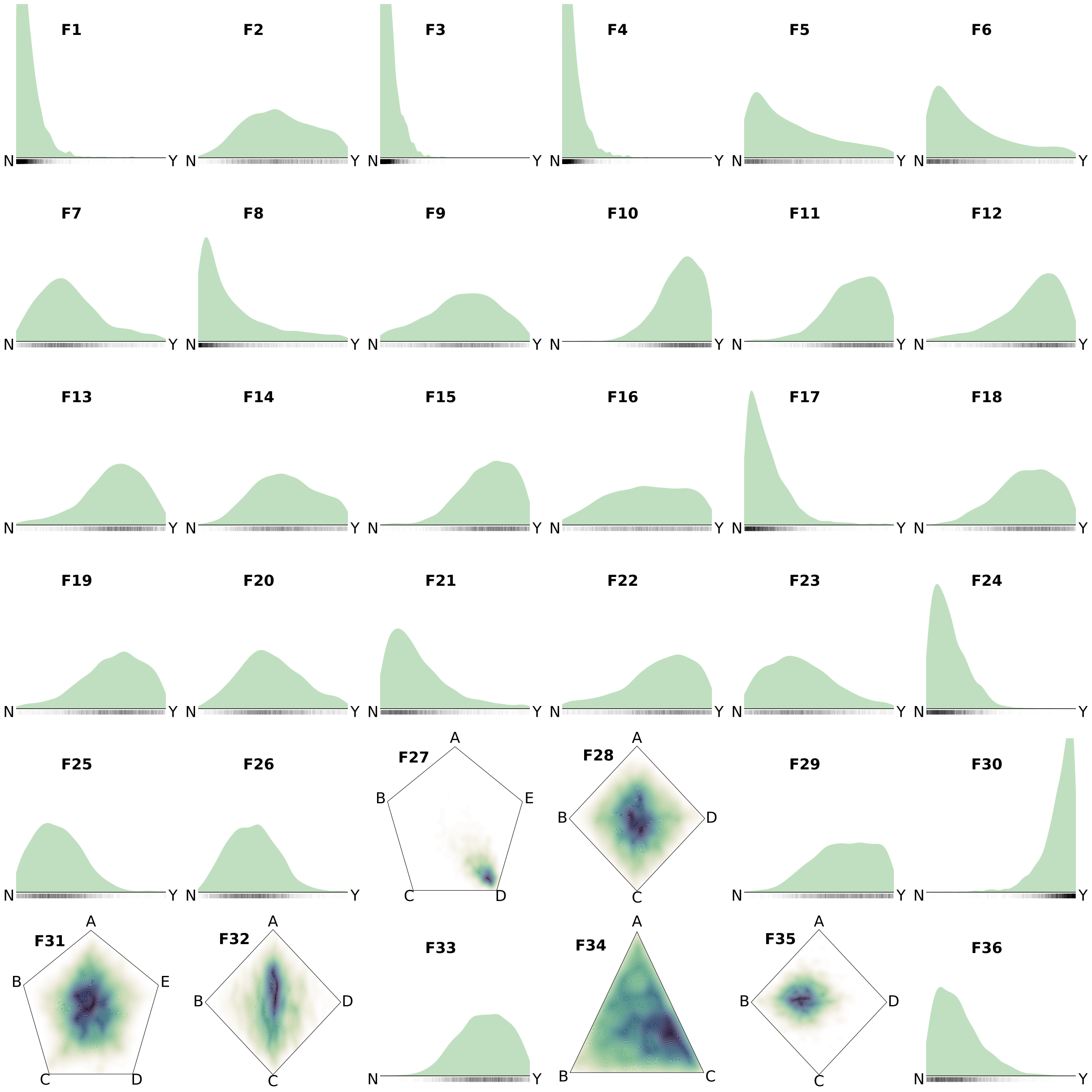


Figure S12: Western South America: areal probabilities in $Z_3$ (the purple contact area in Fig. 5a, Main Paper).

### S9 Case Study: Balkans

For the Balkans case study we coded 47 features for 28 dialects spoken on the Balkans. The feature set consists of three sub-sets:

– Features F1 to F26 comprise linguistic aspects that are not traditionally regarded as constitutive of the Balkans area. The choice is based on two principles. First, the features should be present in dialect varieties and do not necessarily have to be part of the corresponding standard languages (if available, since not all varieties in our sample are standardised). This means that they could have been potentially adapted either by language contact, or by language shift, or because of substrate-adstrate relations. Second, the features need to be present in at least two Balkan dialect varieties.

– Features F27 to F37 comprise a selection of the Standard Average European (SAE) features [Haspelmath, 2001]. Only those SAE features were considered that appear in at least one Balkan variety.

– Features F38 to F47 are traditionally assumed to be constitutive for the Balkan linguistic area [Lindstedt, 2000].

Figure S13 shows the spatial locations of all dialects, Table S3 lists the features used for the analysis. Figure S14 shows that for $n = 1$ all but three languages in the sample are assigned to a single area. Figure S15 plots DIC for models with an increasing number of areas $n$.

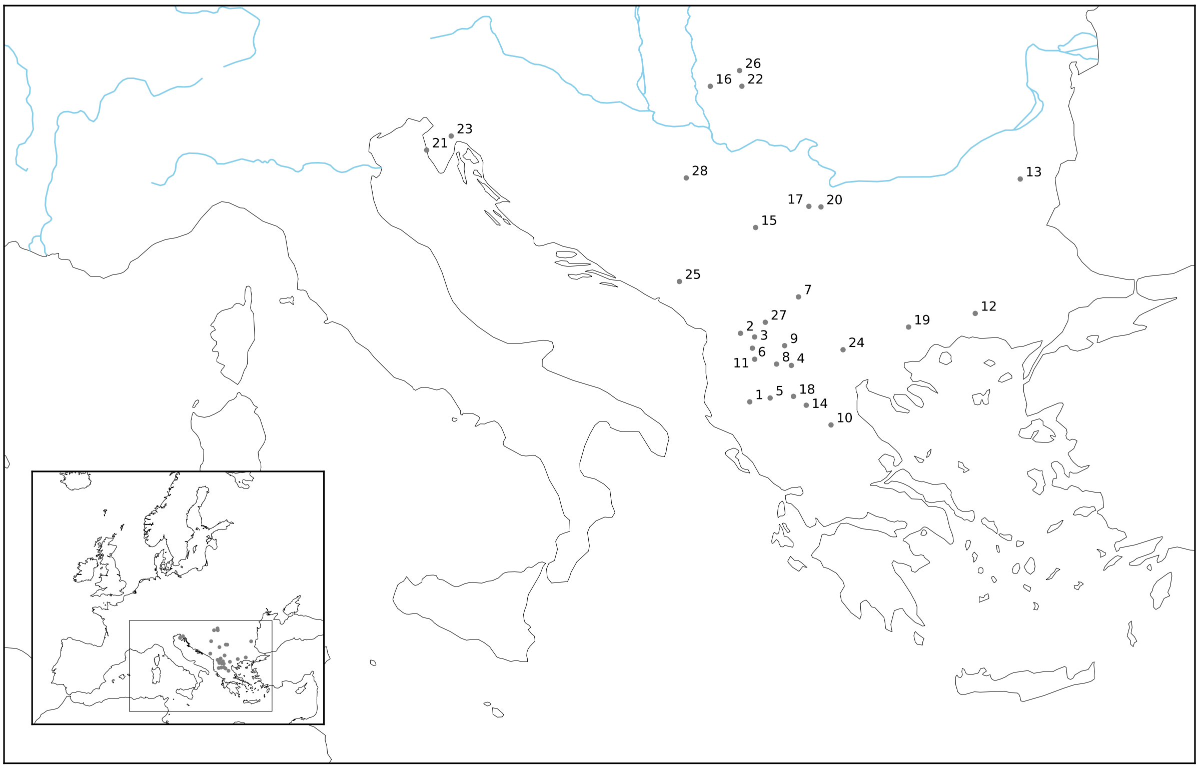

| | | | | | | | | | |
|---|---|---|---|---|---|---|---|---|---|
| 1 | Leshnja | 7 | Kumanovo | 13 | Kaspichan | 19 | Swchalni | 25 | Zavala |
| 2 | Muhhur | 8 | Resen | 14 | Eratyra | 20 | Stakevci | 26 | Vinga |
| 3 | Janche | 9 | Krushevo | 15 | Kobilje | 21 | Porech | 27 | Gostivar |
| 4 | Bitola | 10 | Turia | 16 | Kikinda | 22 | Timishoara | 28 | Osechina |
| 5 | Boboshtica | 11 | Gorna Belica | 17 | Berchinovac | 23 | Zhejan | | |
| 6 | Trebisshte | 12 | Tihomir | 18 | Kostur | 24 | Gevgelija | | |

Figure S13: All languages and dialects coded for the Balkans case study.

Table S3: Features coded in the Balkan case study

| Feature | Description | States |
|---|---|---|
| F1 | Phonemic /ä/ | present, absent |
| F2 | Phonemic /å/ | present, absent |
| F3 | Phonemic /ü/ | present, absent |
| F4 | Phonemic /dz/ | present, absent |
| F5 | Phonemic /h/ | present, absent |
| F6 | Phonemic /$\theta$/ | present, absent |
| F7 | Phonemic palatal nasal | present, absent |
| F8 | Length of vowels | present, absent |
| F9 | Linking articles | present, absent |
| F10 | Mobility of the article within a NP | present, absent |
| F11 | At least one peripheral case (genitive, instrumental, locative, ablative) | present, absent |
| F12 | Inflectional IO marker for M substantives | present, absent |
| F13 | Inflectional IO marker for F substantives | present, absent |
| F14 | Inflectional DO marker for M substantives | present, absent |
| F15 | Inflectional DO marker for F substantives | present, absent |
| F16 | Gender differentiation 3PL personal pronoun used referentially | present, absent |
| F17 | Gender differentiation in 3SF personal pronoun used as verbal clitic in Dat | present, absent |
| F18 | Both volutative and possessive future construction | present, absent |
| F19 | Special form of admirative mood | present, absent |
| F20 | habere-perfect forms | present, absent |
| F21 | esse-perfect forms | present, absent |
| F22 | Admirative mood in the modal system (even if not expressed with a special marker) | present, absent |
| F23 | Perfect as an evidential form | present, absent |
| F24 | Future tense as a habitual form | present, absent |
| F25 | Future-in-the-past as a habitual form | present, absent |
| F26 | Different negation participles for different moods | present, absent |
| F27 | Definite and indefinite articles | present, absent |
| F28 | Relative clauses with relative pronouns | present, absent |
| F29 | 'have'-perfect | present, absent |
| F30 | Dative of external possession | present, absent |
| F31 | Negative pronouns and lack of verbal negation | present, absent |
| F32 | Particles in comparative construction | present, absent |
| F33 | Predominant relative-based equitaties | present, absent |
| F34 | Subject person affixes as strict agreement markers | present, absent |
| F35 | Intensifier-reflexive differentiation | present, absent |
| F36 | Inflectional comparative | present, absent |
| F37 | Presence of // | present, absent |
| F38 | Presence of definite article | present, absent |
| F39 | Definite article (if present) is postpositive | true, false |
| F40 | DO reduplication is regular | true, false |
| F41 | IO reduplication is regular | true, false |
| F42 | Analytic comparative and superlative of adjectives and adverbs | present, absent |
| F43 | Locative model of numerals from 11 to 20 | present, absent |
| F44 | Absence of infinitive | true, false |
| F45 | Volutative future | present, absent |
| F46 | Verb system structure | full (aor., imperf., perf.), perfect-only, absent |
| F47 | Coincidence of directional and spatial interrogative adverb | true, false |

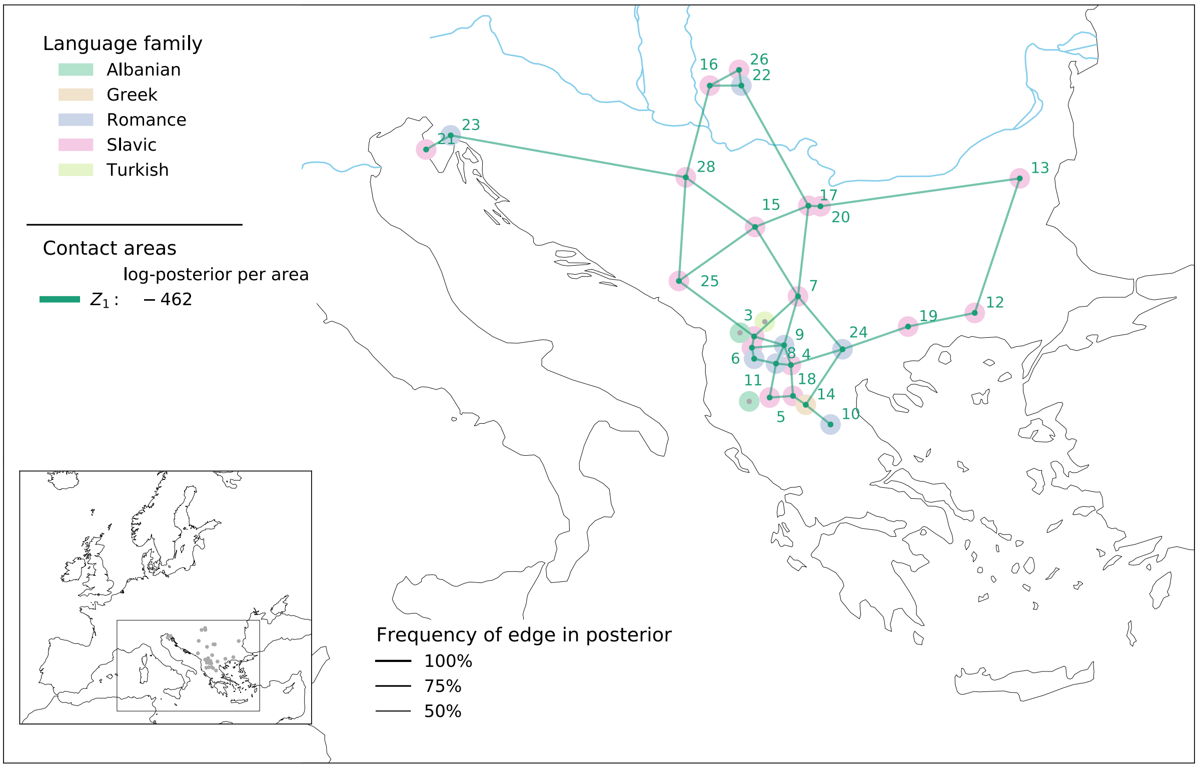


Figure S14: The Balkans as a Sprachbund: For $n = 1$ all but three languages are assigned to a single area $Z_1$.

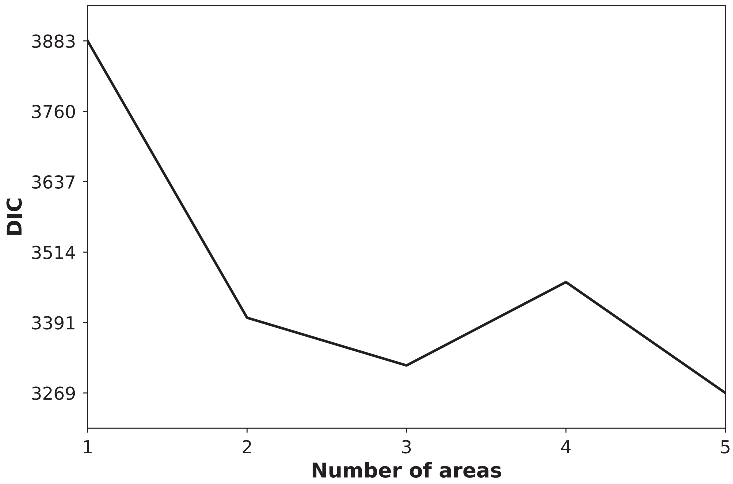


Figure S15: The DIC levels off at $n = 3$, reporting 3 salient contact areas in the Western South American Case study.

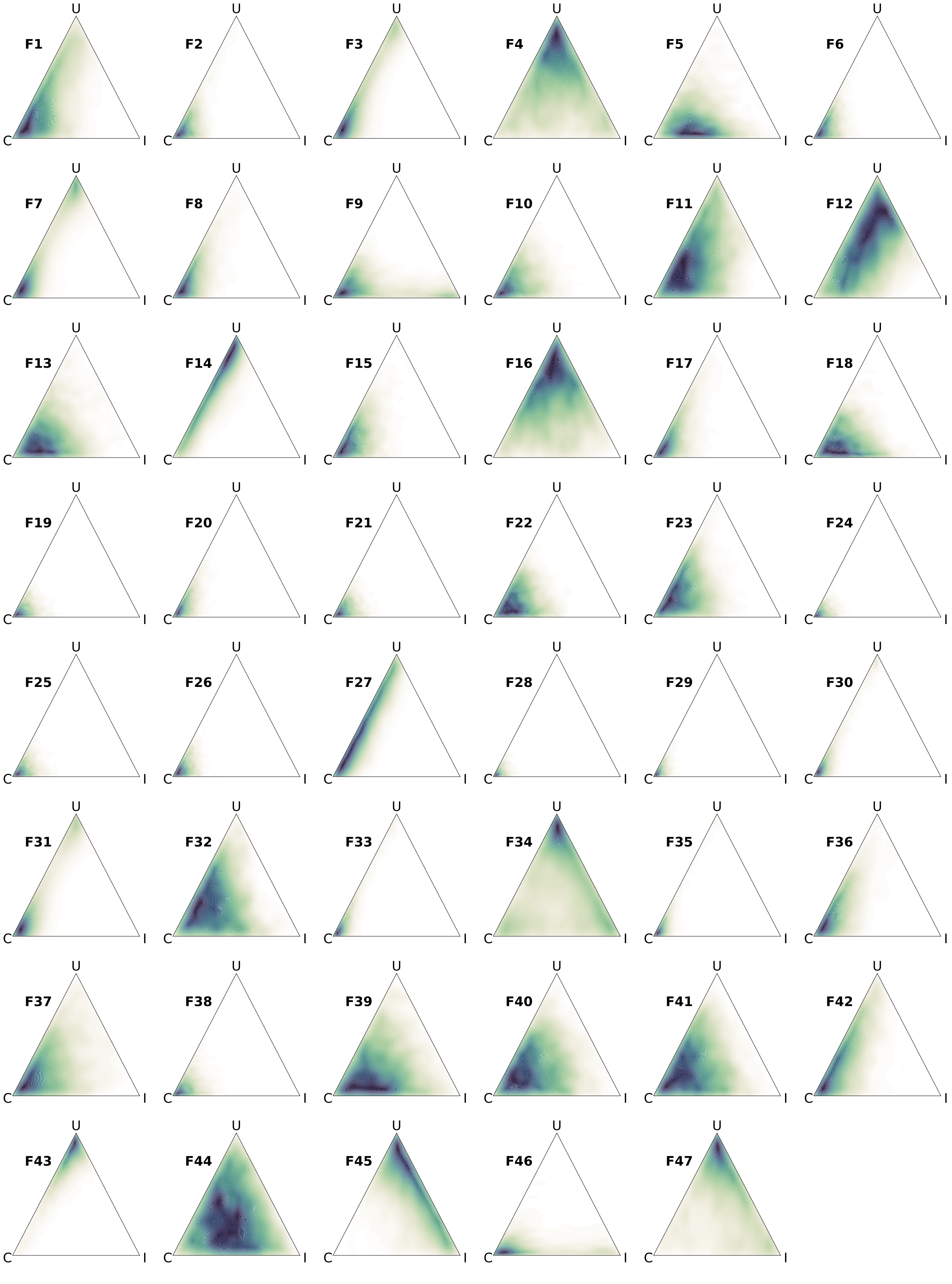


Figure S16: Balkans: weights for all features.

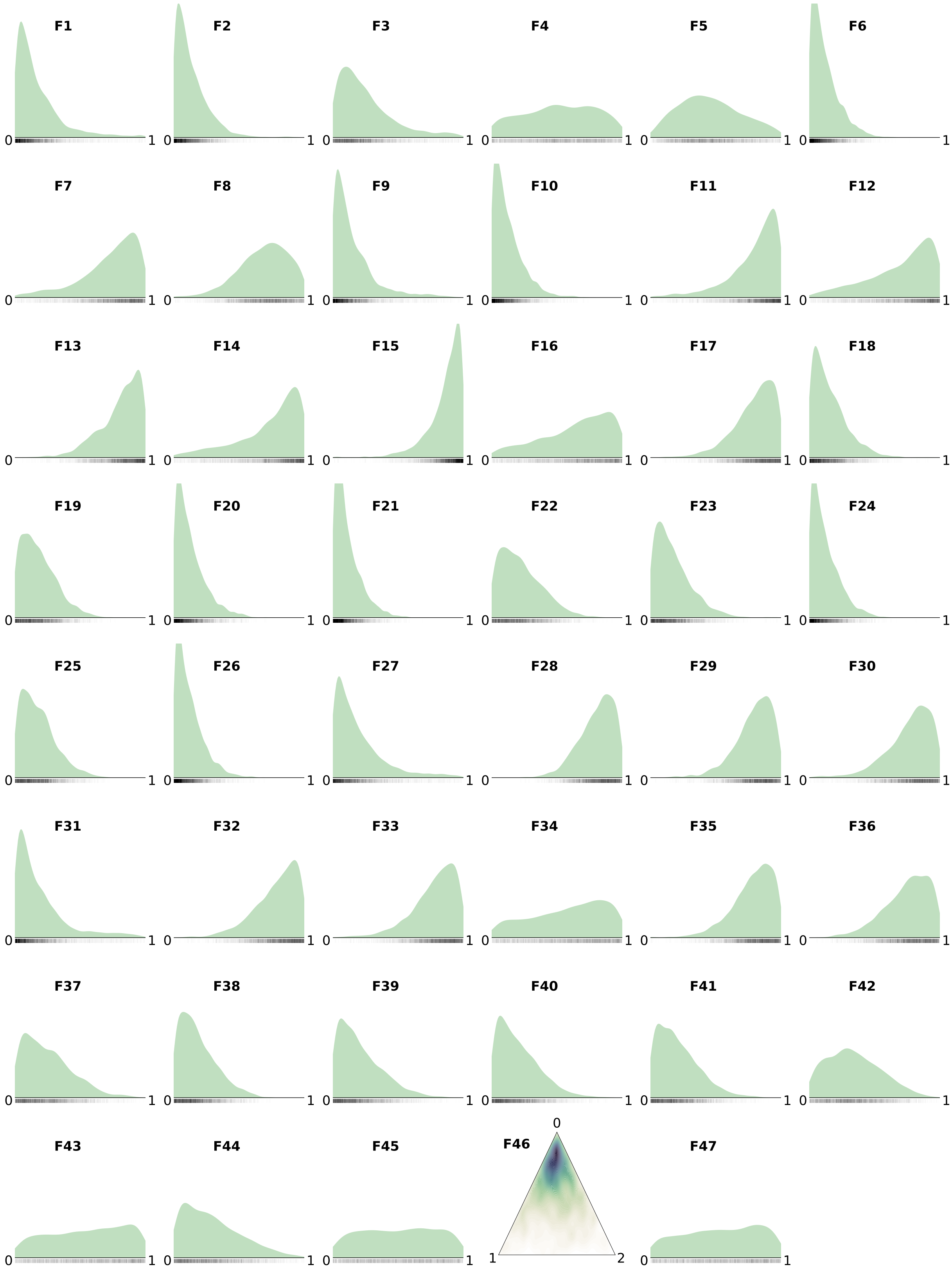


Figure S17: Balkans: areal probabilities in $Z_1$ (the green contact area in Fig. 6a, Main Paper).

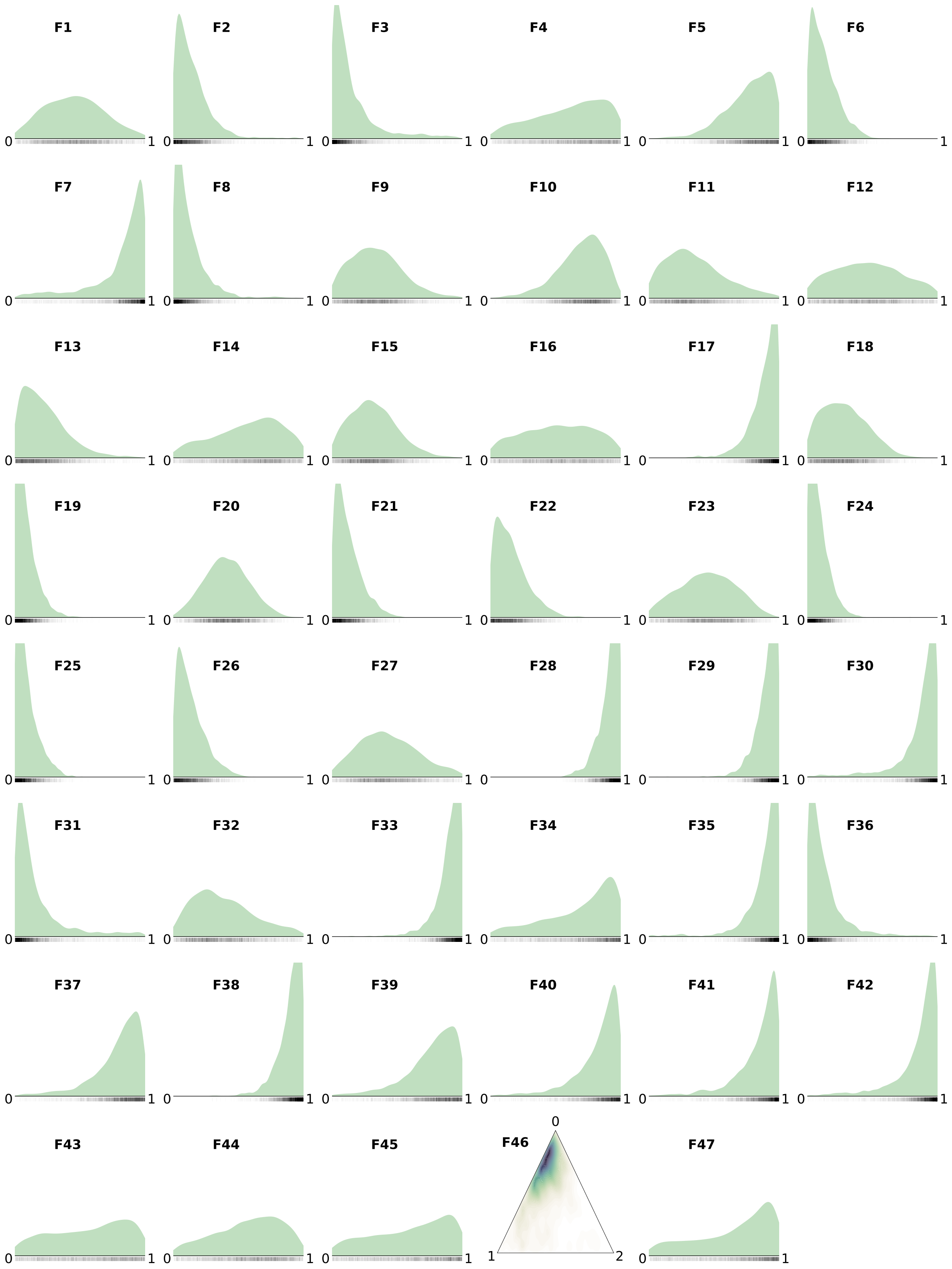


Figure S18: Balkans: areal probabilities in $Z_2$ (the orange contact area in Fig. 6a, Main Paper).

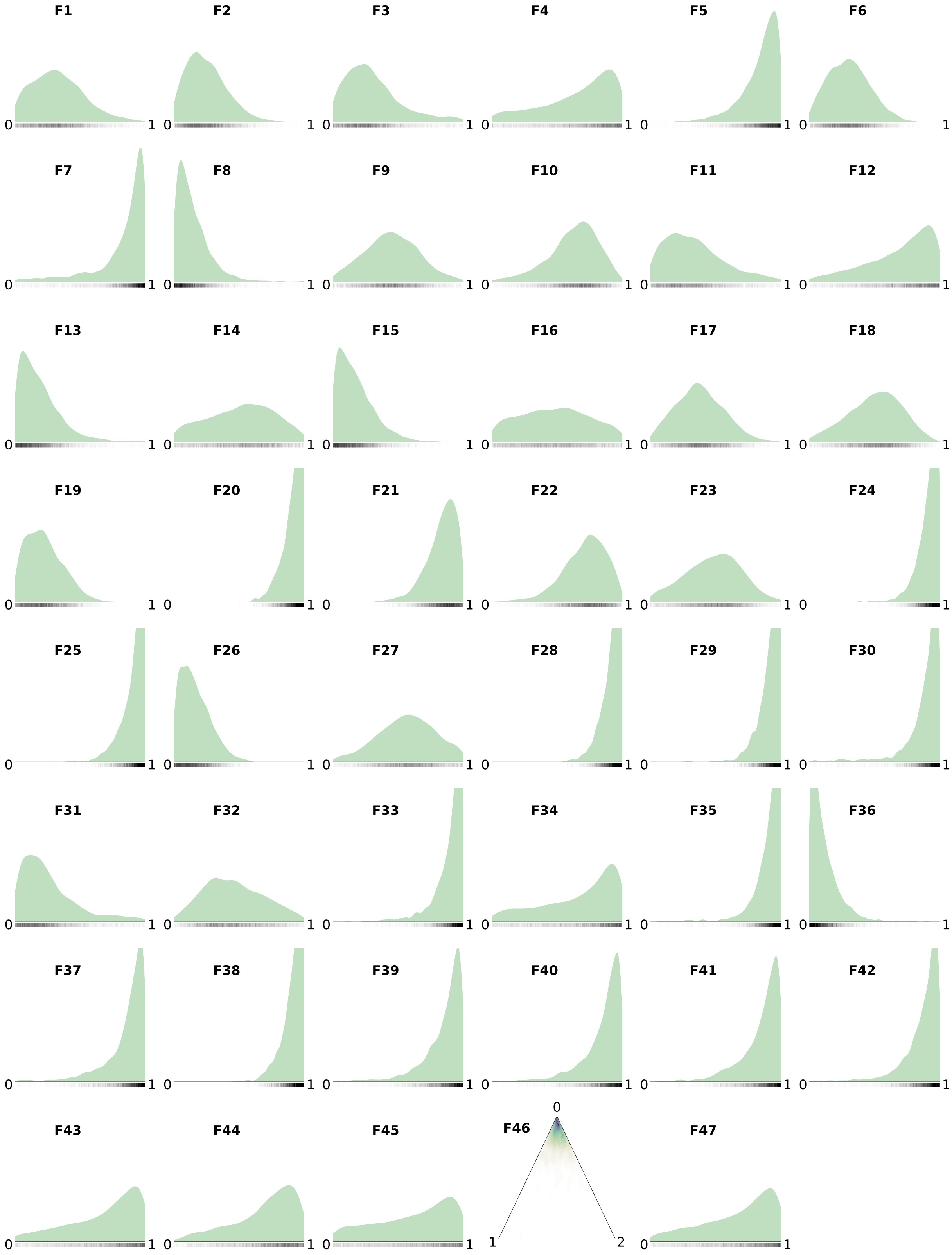


Figure S19: Balkans: areal probabilities in $Z_3$ (the purple contact area in Fig. 6a, Main Paper).

Claire Bowern. The riddle of Tasmanian languages. *Proceedings of the Royal Society of London B: Biological Sciences*, page rspb20121842, 2012.

Claire Bowern. Relatedness as a factor in language contact. *Journal of Language Contact*, 6(2): 411–432, 2013.

Lyle Campbell. Areal linguistics: A closer scrutiny. In Yaron Matras, April McMahon, and Nigel Vincent, editors, *Linguistic areas: Convergence in historical and typological perspective*, pages 1–31. Palgrave MacMillan, Basingstoke, 2006.

Stefan Dedio, Peter Ranacher, and Paul Widmer. Evidence for Britain and Ireland as a linguistic area. *Language*, 95(3):498–522, 2019.

Matthew S. Dryer and Martin Haspelmath, editors. *WALS Online*. Max Planck Institute for Evolutionary Anthropology, Leipzig, 2013. URL `https://wals.info/`.

Michael Dunn, Stephen C Levinson, Eva Lindström, Ger Reesink, and Angela Terrill. Structural phylogeny in historical linguistics: methodological explorations applied in Island Melanesia. *Language*, 84(4):710–759, 2008.

Nick J. Enfield. Areal linguistics and mainland southeast asia. *Annual Revue of Anthropology*, 34:181–206, 2005.

Patience Epps and Lev Michael. *The Areal Linguistics of Amazonia*, page 934–963. Cambridge Handbooks in Language and Linguistics. Cambridge University Press, 2017. doi: 10.1017/9781107279872.033.

Patience Epps, John Huehnergard, and Na'ama Pat-El. Introduction: Contact among genetically related languages. *Journal of Language Contact*, 6(2):209–219, 2013.

Victor A. Friedman. The Balkan Languages and Balkan Linguistics. *Annual Review of Anthropology*, 40(1):275–291, 2011.

Hans Goebl. Le Laboratoire de Dialectométrie de l'Univerisité de Salzbourg. Un bref rapport de recherche. *Zeitschrift für französische Sprache und Literatur*, pages 35–55, 2008.

Wei Hao and John D Storey. Extending tests of hardy–weinberg equilibrium to structured populations. *Genetics*, 213(3):759–770, 2019.

Martin Haspelmath. The european linguistic area: Standard average european. In Martin Haspelmath, Ekkehard König, Wulf Oesterreicher, and Wolfgang Raible, editors, *Language typology and language universals. (Handbücher zur Sprach- und Kommunikationswissenschaft)*, page 1492–1510. de Gruyter, Berlin, 2001.

Bernd Heine. *A linguistic geography of Africa*. Cambridge University Press, 2008.

Péter Jeszenszky, Philipp Stoeckle, Elvira Glaser, and Robert Weibel. A gradient perspective on modeling interdialectal transitions. *Journal of Linguistic Geography*, 6(2):78–99, 2018. doi: 10.1017/jlg.2019.1.

Olga Krasnoukhova. *The Noun Phrase in the Languages of South America*. PhD thesis, Radboud Universiteit Nijmegen, 2012.

Jouko Lindstedt. Linguistic balkanization: contact-induced change by mutual reinforcement. *Studies in Slavic and General Linguistics*, 28:231–246, 2000.

Jan Lundberg. Classifying dialects using cluster analysis. *Master's thesis, Göteborg University*, 2005.

Colin Masica. The definition and significance of linguistic areas: methods, pitfalls, and possibilities (with special reference to the validity of south asia as a linguistic area). In Peri Bhaskararao and Karumuri Venkata Subbarao, editors, *Tokyo Symposium on South Asian languages: contact, convergence, and typology*, pages 205–267. Sage Publications, New Delhi, 2001.

Luisa Miceli and Alan Dench. *The Areal Linguistics of Australia*, page 732–757. Cambridge Handbooks in Language and Linguistics. Cambridge University Press, Cambridge, 2017.

Pieter Muysken. Conceptual and methodological issues in areal linguistics. In Pieter Muysken, editor, *From Linguistic Areas to Areal Linguistics*, pages 1–23. John Benjamins, 2008.

Pieter Muysken, Harald Hammarström, Joshua Birchall, Rik Van Gijn, Olga Krasnoukhova, and Neele Müller. Linguistic areas: bottom-up or top-down? the case of the guaporé-mamoré. In Bernard Comrie and Lucia Golluscio, editors, *Language Contact and Documentation / Contacto lingüístico y documentación*, pages 205–238. Walter de Gruyter, Berlin, 2014.

Michael Noonan. *Genetic Classification and Language Contact*, chapter 2, pages 48–65. John Wiley & Sons, Ltd, Malden and Oxford, 2010.

Jonathan K Pritchard, Matthew Stephens, and Peter Donnelly. Inference of population structure using multilocus genotype data. *Genetics*, 155(2):945–959, 2000.

Andrew Rambaut, Alexei J Drummond, Dong Xie, Guy Baele, and Marc A Suchard. Posterior summarization in Bayesian phylogenetics using Tracer 1.7. *Systematic Biology*, 67(5):901, 2018.

Ger Reesink and Michael Dunn. Systematic typological comparison as a tool for investigating language history. *Language Documentation & Conservation*, 5:34–71, 2012.

Ger Reesink, Ruth Singer, and Michael Dunn. Explaining the linguistic diversity of Sahul using population models. *PLoS biology*, 7(11):e1000241, 2009.

Yves Scherrer and Philipp Stoeckle. A quantitative approach to Swiss German–dialectometric analyses and comparisons of linguistic levels. *Dialectologia et Geolinguistica*, 24(1):92–125, 2016.

Thomas Stolz. No sprachbund beyond this line. on the age-old discussion of how to define a linguistic area. In Paolo Ramat and Thomas Stolz, editors, *Mediterranean languages. Papers from the MEDTYP workshop, Tirrenia, June 2000*, pages 259—-281. Universitätsverlag Dr. N. Brockmeyer, Bochum, 2002.

Thomas Stolz. All or nothing. In Yaron Matras, April McMahon, and Nigel Vincent, editors, *Linguistic areas: Convergence in historical and typological perspective*, pages 32–50. Palgrave MacMillan, Basingstoke, 2006.

Kaj Syrjänen, Terhi Honkola, Jyri Lehtinen, Antti Leino, and Outi Vesakoski. Applying Population Genetic Approaches within Languages. *Language Dynamics and Change*, 6(2):235–283, 2016.

Benedikt Szmrecsanyi. Corpus-based dialectometry: a methodological sketch. *Corpora*, 6(1): 45–76, 2011.

Rik van Gijn. The Andean foothills and adjacent Amazonian fringe. In Loretta O'Connor and Pieter Muysken, editors, *The Native Languages of South America. Origins, Development, Typology*, pages 102–125. Cambridge University Press, Cambridge, 2014.

Rik van Gijn. Separating layers of information. *Advances in Contact Linguistics: In honour of Pieter Muysken*, 57:161, 2020. doi: 10.1075/coll.57.05van.

Rik Van Gijn and Max Wahlström. Linguistic areas. In Rik van Gijn, Max Wahlström, Hanna Ruch, and Anja Hasse, editors, *Language contact: bridging the gap between individual interactions and areal patterns*. forthcoming.

Martijn Wieling and John Nerbonne. Bipartite spectral graph partitioning for clustering dialect varieties and detecting their linguistic features. *Computer Speech & Language*, 25(3):700–715, 2011.
